# Changes in Histone Isoform Abundnance and Histone Post-Translational Modifications during Anoxia Tolerance and Recovery in WS40NE cells of *Austrofundulus limnaeus*

**DOI:** 10.1101/2025.09.04.674156

**Authors:** Chelsea Hughes, Elizabeth A. Mojica, Dietmar Kültz, Jason E. Podrabsky

**Affiliations:** Department of Biology, Center for Life in Extreme Environments, Portland State University, P.O. Box 751, Portland, OR 97201, USA; Nicholas School of the Environment, Duke University, 308 Research Dr, Levine Science Research Center, Durham, NC 27708, USA; Department of Animal Sciences and Genome Center, University of California, Davis, One Shields Avenue, Meyer Hall, Davis, CA 95616, USA; Department of Integrative Biology, Oregon State University, 2701 SW Campus Way, Corvallis, OR 97331, USA

**Keywords:** mass spectrometry, epigenetics, histone post-translational modifications, anoxia, stress tolerance

## Abstract

Anoxia is an often-lethal stressor to vertebrates, yet some vertebrates have adapted cellular mechanisms to survive in anoxic conditions. Embryos of the annual killifish *Austrofundulus limnaeus* have the greatest tolerance to anoxia of all vertebrates, yet the epigenetic mechanisms that support their anoxia tolerance are unknown. Using mass spectrometry, 1043 unique biologically relevant histone post-translational modifications (unimod+histone residue) were detected in WS40NE cells, representing thirteen types of biologically relevant histone post-translational modifications (hPTMs) present during normoxia, 1 d anoxia, 4 d anoxia, and aerobic recovery from anoxia. Of these 1043 hPTMs, 816 were significantly differentially expressed in at least one comparison. Thirty-six significant hPTMs were considered highly condition dependent. Additionally, at least four histone isoforms were differentially expressed, representing H2A, H2B, and H3 isoforms. Our data suggests that specific histone modifications as well as changes in histone isoform abundance in WS40NE cells may be necessary to successfully respond to extreme changes in oxygen availability.

## Introduction

All organisms must respond to their environment, which offers both opportunities and threats. By changing which genes are expressed, organisms can modify their phenotype to take advantage of opportunities and neutralize threats. However, environmental stressors can often lead to aberrant gene expression in organisms that lack these adaptations. This is particularly true during anoxic stress in mammals, which can cause apoptosis and inappropriate reentry into the cell cycle in mammalian cells [1, 2]. Specifically in mammalian neurons, apoptosis is associated with cellular dysregulation that leads these terminally differentiated cells to re-enter the cell cycle, which often leads to catastrophic cell failure [2]. Additionally, mitochondrial dysregulation is associated with anoxia due to inhibition of the electron transport chain (ETC) and depolarization of the inner mitochondrial membrane that can lead to apoptosis [3–5]. The subsequent cell death causes tissue damage, which continues when oxygen is returned to the affected area, a phenomenon known as ischemia-reperfusion injury in mammals [6]. Ischemia-reperfusion increases reactive oxygen species (ROS) such as the superoxide anion (O^−2^), hydrogen peroxide (H_2_O_2_), and hydroxyl radical (OH^−^) [7]. While low levels of ROS are a normal byproduct of mitochondrial metabolism [8], high levels of ROS can lead to neuronal death within 5 minutes [9, 10]. In contrast, a handful of vertebrates have adapted mechanisms to modify their gene expression to survive environmental stressors, including osmotic, thermal, ROS, and anoxia with little or no damage [11–19].Therefore, there is a need to understand how gene expression is regulated in anoxia tolerant vertebrates to understand how they avoid these fates. This study focuses on epigenetic histone modifications in response to anoxia in a cell line isolated from anoxia-tolerant embryos of the annual killifish, *Austrofundulus limnaeus*.

Epigenetic responses to a variety of stressors are documented in multiple organisms [20–23]. However, few histone post-translational modifications (hPTMs) have been investigated in response to hypoxia or anoxia. Additionally, these data often come from anoxia intolerant organisms that are experiencing a disease state and are likely failing [24, 25]. Additionally, this research often focuses only on well-known residues and specific types of hPTMs such as methylation [26, 27]. For example, histone-3 lysine-4 trimethylation (H3K4me3) and H3K36me3 are known to change in HeLa cells within 1 h of hypoxic exposure, but HeLa cells begin to die within 1 d of severe hypoxia [28, 29]. Limited research exists on histone modifications in anoxia tolerant organisms, but this work again focuses on well-known modifications such as acetylation and methylation [30]. It is unknown how less studied hPTMs, such as lactylation, 4-hydroxynonenal, and oxidation are impacted by anoxic stress, particularly in anoxia tolerant species. Even better characterized hPTMs like ubiquitylation are not well understood in anoxia. While ubiquitylation of the proteome has been studied in ischemia and anoxia recovery [31, 32], to our knowledge, anoxia-induced histone ubiquitylation has never been explored in a continuous cell line.

Embryos of the annual killifish *Austrofundulus limnaeus* have the greatest tolerance to anoxia of all vertebrates, making them the ideal model to study the cellular mechanisms necessary for anoxia tolerance. This remarkable tolerance of anoxia is supported by the ability of embryos to arrest development and enter a state of anoxia-induced quiescence [33]. The global histone landscape of developing embryos (Wourms’ Stage 36) under normoxia and after 24 h of anoxia suggests a stabilization of both histone isoform and hPTM abundance [34]. This is surprising, as hPTMs are known to respond to stressors quickly [22, 28, 35, 36]. However, this stabilization represents an averaged histone landscape for multiple cell types [34]. It is unclear how specific cell lines of *A. limnaeus* respond to anoxia, particularly in the absence of tissue-dependent cell signaling. Additionally, no research has investigated how the histone landscape of *A. limnaeus* changes when exposed to longer bouts of anoxia or during recovery from anoxia.

PSU-AL-WS40NE (from now on WS40NE) is a neuroepithelial cell line that was isolated from an embryonic (Wourms’ Stage 40) *A. limnaeus* tissue explant. These cells can survive for at least 49 days without oxygen [37], while SH-SY5Y human neuroblastoma cells can survive anoxia for only approximately 1.3 days [38], and mouse N2A cells can survive for just over 3 days [37]. Here, we describe for the first time the global histone landscape of WS40NE cells exposed to anoxia and recovery from anoxia. We hypothesize cells undergo extensive changes to histone isoforms and the hPTM landscape in response to anoxia and aerobic recovery from anoxia.

## Results

### Histone Abundance

Our methods resulted in the identification of 1,927 peptides across at least twenty-three distinct histone proteins based on their unique protein accession number. These proteins were representative of H1, H2A, H2B, and H3 isoforms. No H4 isoforms were detected. Sequence coverage ranged from 17-100% (Fig. 1). At least four histones were differentially expressed across the four experimental time points (Fig. 1).

**Figure 1.**
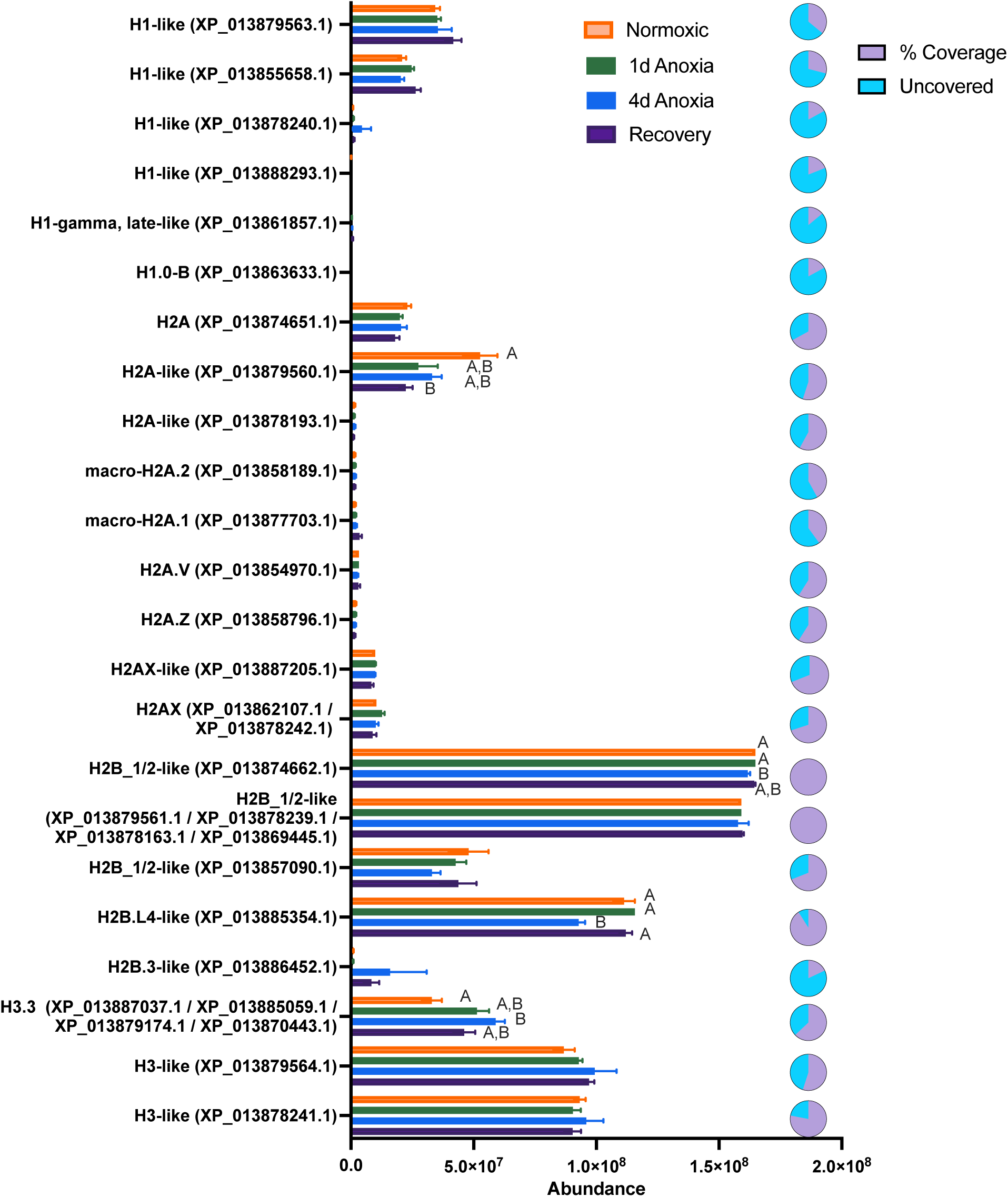
Histone isoform abundance in WS40NE cells. Twenty distinct histone proteins were identified based on their unique protein accession number, as well as three groups of histones that could not be distinguished. Bar graphs represent the quantile normalized mean ± SEM of histone relative abundance during normoxia (orange), 1 d anoxia (green), 4 d anoxia (blue), and 1 d of recovery (dark purple). Letters over bars represent statistically significant isoform abundances determined by pairwise comparisons. Pie charts represent the sequence coverage (light purple) and the percent of peptide sequences that were not detected (blue) for each isoform. Sequence coverage is defined as the total observed protein sequence length divided by the total protein length. Note that due to different peptide ionization efficiencies, histone isoform abundance can only be compared across conditions and not to one another within a condition.

Histone H3.3 protein increased significantly within 4 d of anoxia, while an H2A-like protein decreased during anoxia compared to normoxia, decreasing significantly during recovery. Two histone H2B isoforms decreased in abundance after 4 d of anoxia. However, principal component analysis demonstrated broad sample similarity across conditions when considering the abundances of all histone isoforms (Fig. 2).

**Figure 2.**
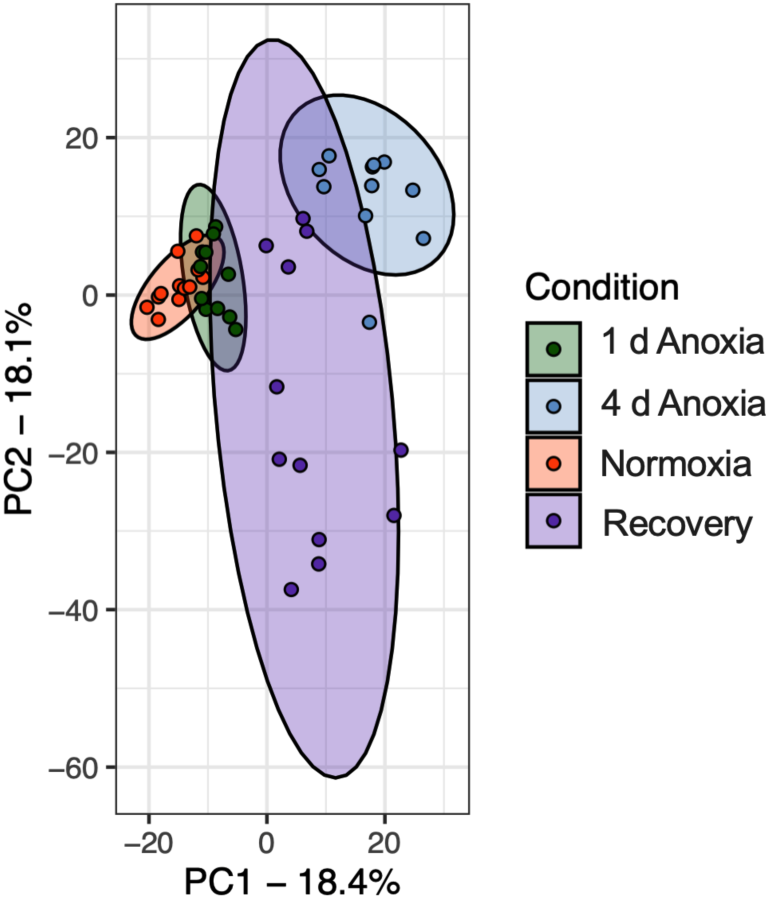
Principal component analysis of histone isoform abundance in WS40NE cells exposed to anoxia and aerobic recovery from anoxia. The mean quantile normalized abundance values of all histone isoforms were used to construct a PCA Plot. Normoxic (purple), 1 d anoxic (green), 4 d anoxic (blue), and recovery samples (pale blue) are plotted according to PC1 and PC2 with a 95% confidence ellipse surrounding each cluster.

### Histone Post-translational Modifications

Post-translational modifications were evaluated in a residue-specific manner to determine specific sites of differential modification on each histone. Principal component analysis (PCA) illustrates that, globally, the hPTM landscape is most unique after 4 d of anoxia compared to normoxia (Fig. 3C). It is worth noting the high variation in PC2 for the recovery time point (Fig. 3C). Out of 1293 unique hPTMs detected, 1043 unique hPTMs (hPTM type+residue) were considered biologically relevant, representing thirteen types of biologically relevant PTMs (Fig. 3A,B). Four hundred and sixty-one residues were modified with biologically relevant hPTMs, with many residues being modified with multiple types of hPTMs. These types of modifications included commonly studied modifications such as acetylation, methylation, phosphorylation, and ubiquitylation, and less studied modifications such as lactylation and 4-hydroxynonenal. All thirteen types of modifications were statistically significant on at least one residue in at least one comparison, with oxidation/hydroxylation and dioxidation having the greatest number of significantly modified residues, followed closely by dehydration and phosphorylation (Fig. 3A). As expected, the percent sequence coverage is positively correlated with the number of significant hPTMs (Fig. 3D, r = 0.7, p = 0.0003), and thus for certain histones our ability to draw global conclusions is somewhat limited. However, several isoforms, such as H1-like (XP_013879563.1), do not follow this trend, suggesting these are highly modified isoforms (Fig. 3D). Of the unique biologically relevant PTMs detected, 816 were significant in at least one comparison (Fig. 3E). Hundreds of significant differential hPTMs were present across all four timepoints with 4 d of anoxia having the highest number compared to other time points (Fig. 3E).

**Figure 3.**
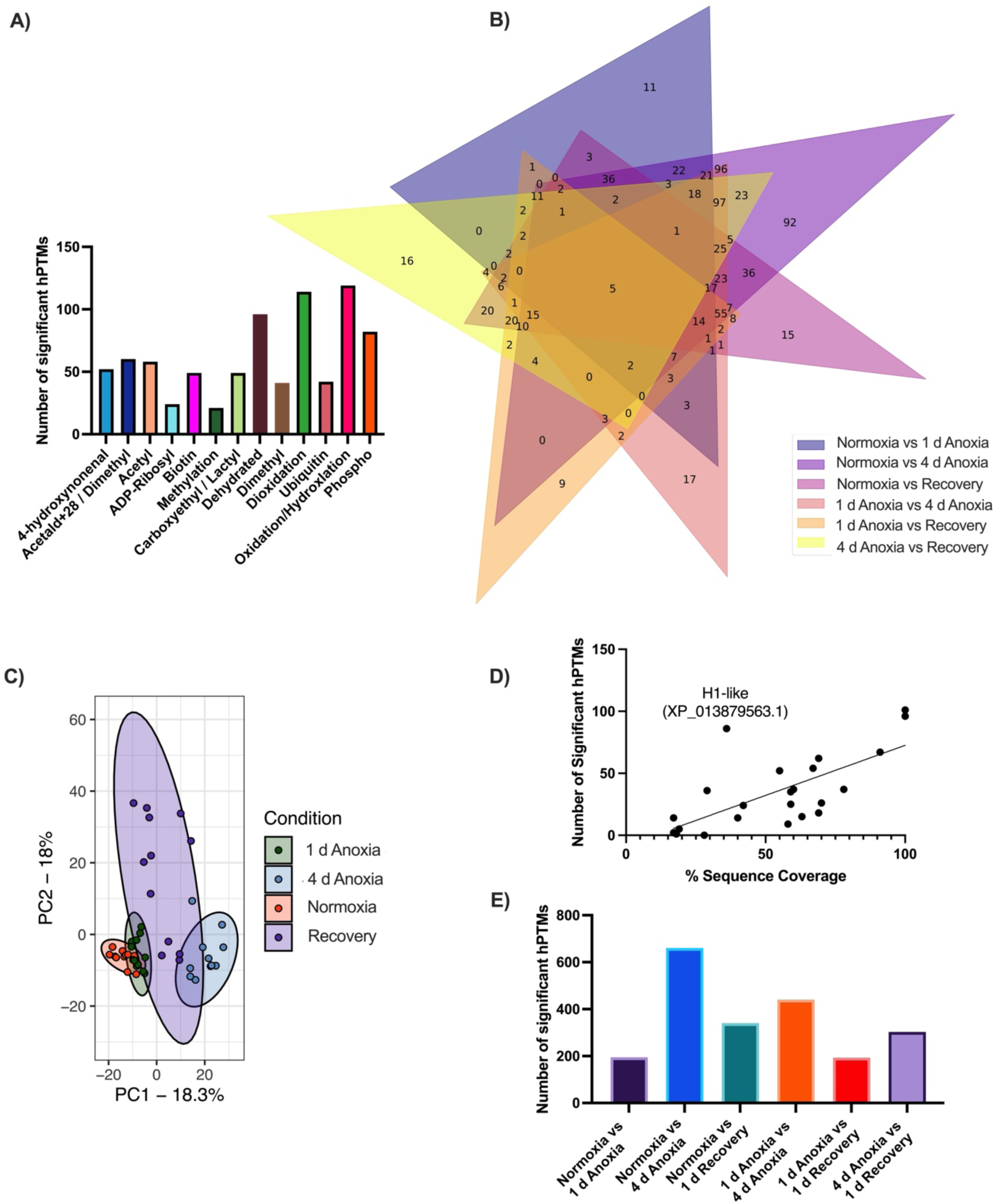
Quantification of global patterns of significant histone post-translational modifications in WS40NE cells exposed to anoxia and recovery from anoxia. **(A)** The number of hPTMs with statistically significant changes in abundance were plotted by modification type across all histone isoforms**. (B)** Venn diagram illustrating the occurrence of the 297 unique and biologically-relevant hPTMs across the experimental conditions used in this study. The Venn diagram is schematic and non-proportional. The numbers within each colored section indicate the number of significantly differentially expressed histone modifications shared among the indicated pairwise comparisons. **(C)** PCA plot of the M-values of all hPTMs indicate that hPTM patterns at 4 d of anoxia are unique compared to the other experimental conditions. It is also noteworthy that variation in global patterns of hPTMs appears to be high during aerobic recovery. Individual samples (data points) are surrounded by 95% confidence ellipses for each treatment group. **(D)** There is a positive correlation (r = 0.7, p = 0.0003) between percent sequence coverage for each histone isoform and the number of significant hPTMs identified. However, several isoforms do not follow this trend, suggesting these are highly modified isoforms. **(E)** The bar graph represents the number of significantly expressed hPTMs in each t-test comparison.

Thirty hPTMs were highly abundant (having an average relative abundance of 50% or greater) across all conditions. Of the high abundance hPTMs, 15 displayed an average relative abundance across all samples of exactly 100% during all conditions, meaning that each specific residue was always modified by that PTM (Table 1). Five histones had at least one hPTM site with 100% relative abundance, representing H1, H2A, and H3 histones. At least 8 distinct types of PTMs are represented in this group with dehydration accounting for 30% of these stable modifications.

**Table 1.** Histone post-translational modifications with 100% relative abundance in WS40NE cells. Fifteen hPTMs, across 5 histone isoforms, always occurred at the same residue across all experimental conditions. At least eight distinct types of PTMs are represented.

| Protein Accession | Protein Description | Amino Acid +<br>Position | PTM Description |
| --- | --- | --- | --- |
| XP_013861857.1 | H1-gamma,_late-like | 196K | Acetylation |
| XP_013861857.1 | H1-gamma,_late-like | 175K | Biotinylation |
| XP_013861857.1 | H1-gamma,_late-like | 173P | Oxidation/Hydroxylation |
| XP_013861857.1 | H1-gamma,_late-like | 191K | Acetylation |
| XP_013861857.1 | H1-gamma,_late-like | 157K | Acetylation |
| XP_013861857.1 | H1-gamma,_late-like | 228K | Dioxidation |
| XP_013861857.1 | H1-gamma,_late-like | 24R | 4-hydroxynonenal |
| XP_013861857.1 | H1-gamma,_late-like | 46S | Phosphorylation |
| XP_013878240.1 | H1-like | 64S | Dehydrated |
| XP_013885354.1 | H2B.L4-like | 46S | Dehydrated |
| XP_013885354.1 | H2B.L4-like | 55S | Dehydrated |
| XP_013885354.1 | H2B.L4-like | 43T | Dehydrated |
| XP_013885354.1 | H2B.L4-like | 31Y | Dehydrated |
| XP_013878241.1 | H3-like | 24R | Oxidation/Hydroxylation |
| XP_013878241.1 | H3-like | 17K | Oxidation/Hydroxylation |

Individual hPTMs at specific residues were also evaluated, and the most condition-dependent hPTMs were identified. Significant differential modifications were detected on all groups of histones (H1, H2A, H2B, H3), with H1-gamma late-like (XP_013861857.1) being the only isoform that did not have any significant hPTMs detected. Thirty-seven hPTMs were considered highly condition dependent, being significantly differentially expressed across 5-6 comparisons (Table 2).

**Table 2.**
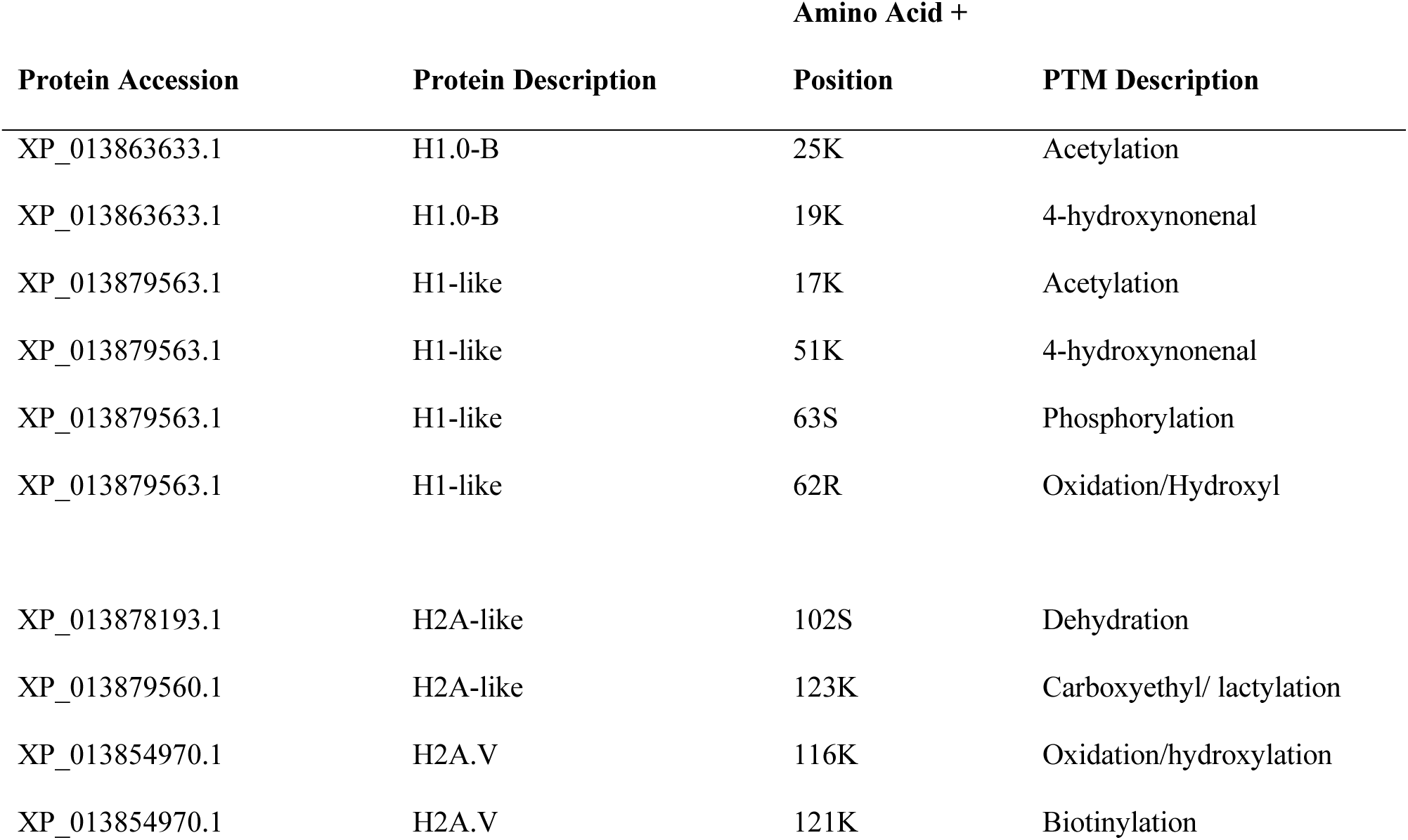
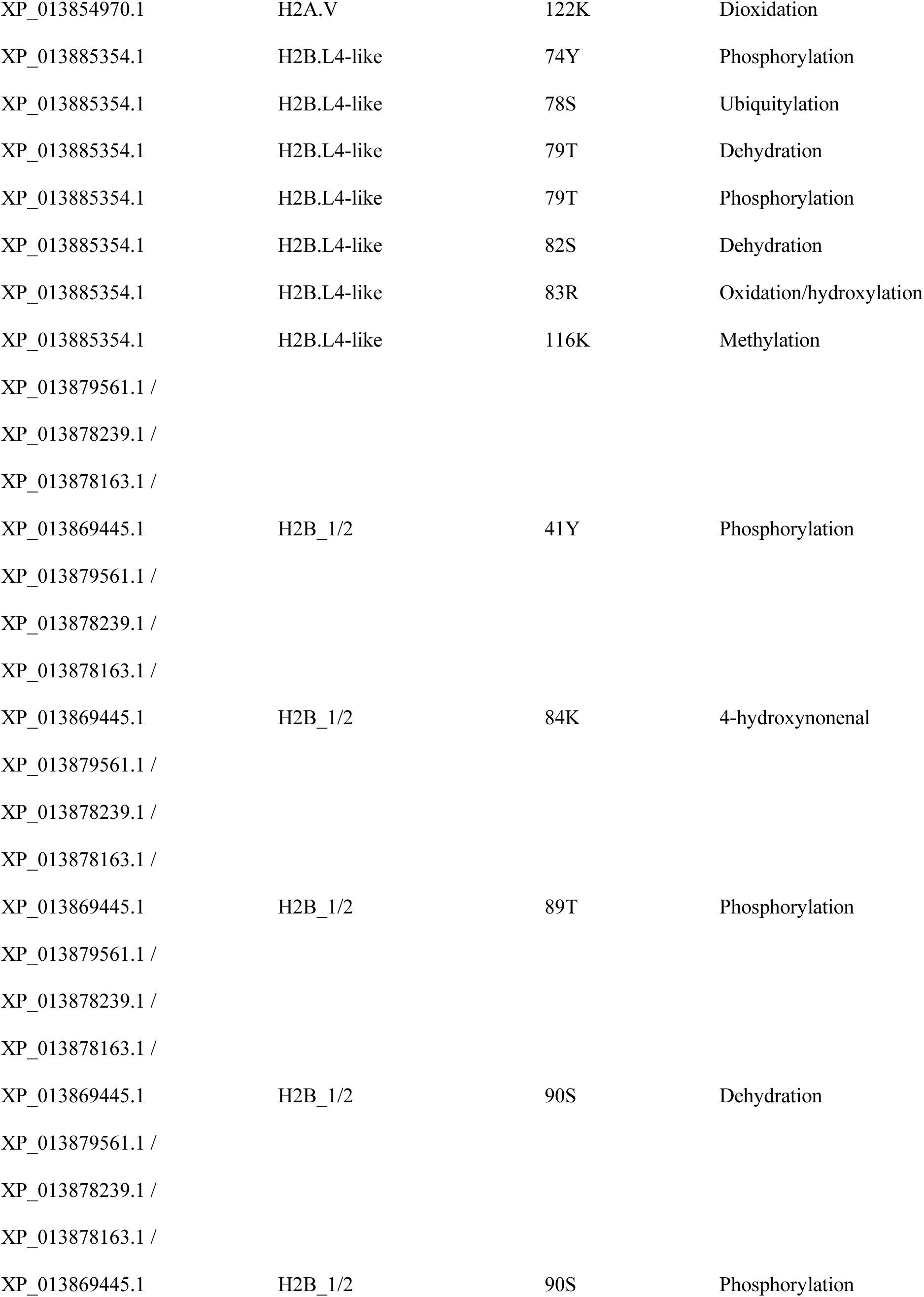
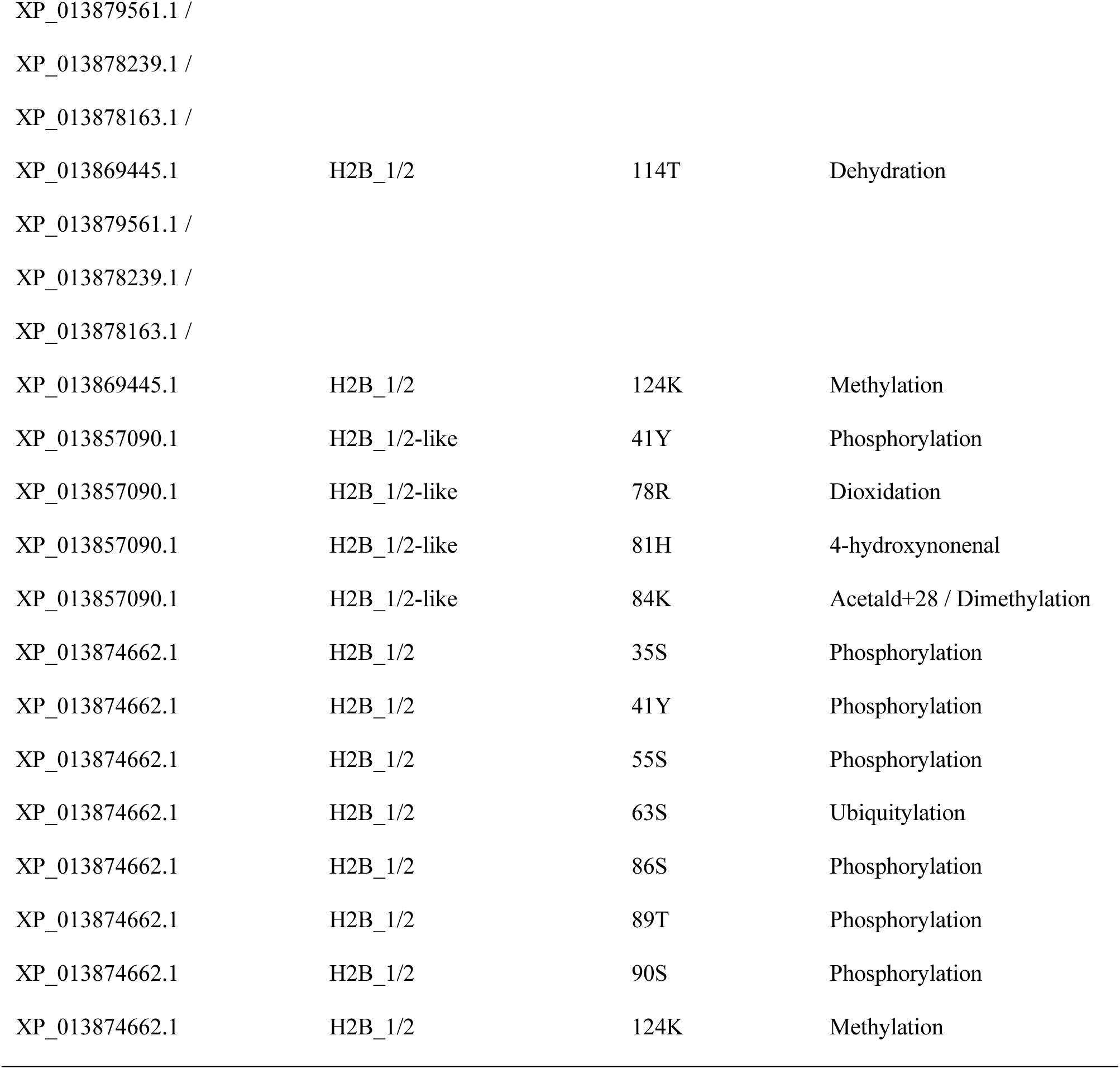
Highly condition-dependent histone post-translational modifications in WS40NE cells exposed to anoxia and aerobic recovery from anoxia. To be considered highly condition-dependent, hPTMs had to be significant across at least 5 of 6 different comparisons. Thirty-seven hPTMs were selected as the most condition dependent. Differential modifications of histone residues were evaluated using a two-tailed t-test with the Benjamini-Hochberg correction applied to account for multiple hypothesis testing.

Globally, hPTMs did not cover the majority of the residues that are theoretically available. Lysine residues were the most modified with 48.7-51.4% of the available residues modified across the different experimental conditions (Fig. 4A). The modification of specific amino acid residues varied based on experimental conditions (Fig. 4A). Lysines had the most types of modifications, with 10 distinct hPTMs detected globally, and of these, oxidation/hydroxylation, dioxidation, and acetylation were the most common (Fig. 4A). Serine (S) had the second most types of modifications, with 5 distinct hPTMs occurring globally; dehydration and phosphorylation dominated serine modifications (Fig. 4A). Globally, there were significant changes in all types of hPTMs across all six comparisons (Fig. 4B,C): (a) 5 comparing normoxia to 24 h anoxia, (b) 12 comparing normoxia to 4 d anoxia, (c) 8 comparing normoxia to 24 h recovery,(d) 6 comparing 24 h anoxia to 4 d anoxia, (e) 4 comparing 24 h anoxia to 24 h recovery, and (f) 2 comparing 4 d anoxia to 24 h recovery. The majority of the most common modifications decreased significantly during anoxia and remained low even after 24 h of aerobic recovery (Fig. 4B,C). Notably, acetylation, dehydration, oxidation, and dioxidation all increase during anoxia. (Fig. 4B,C).

**Figure 4.**
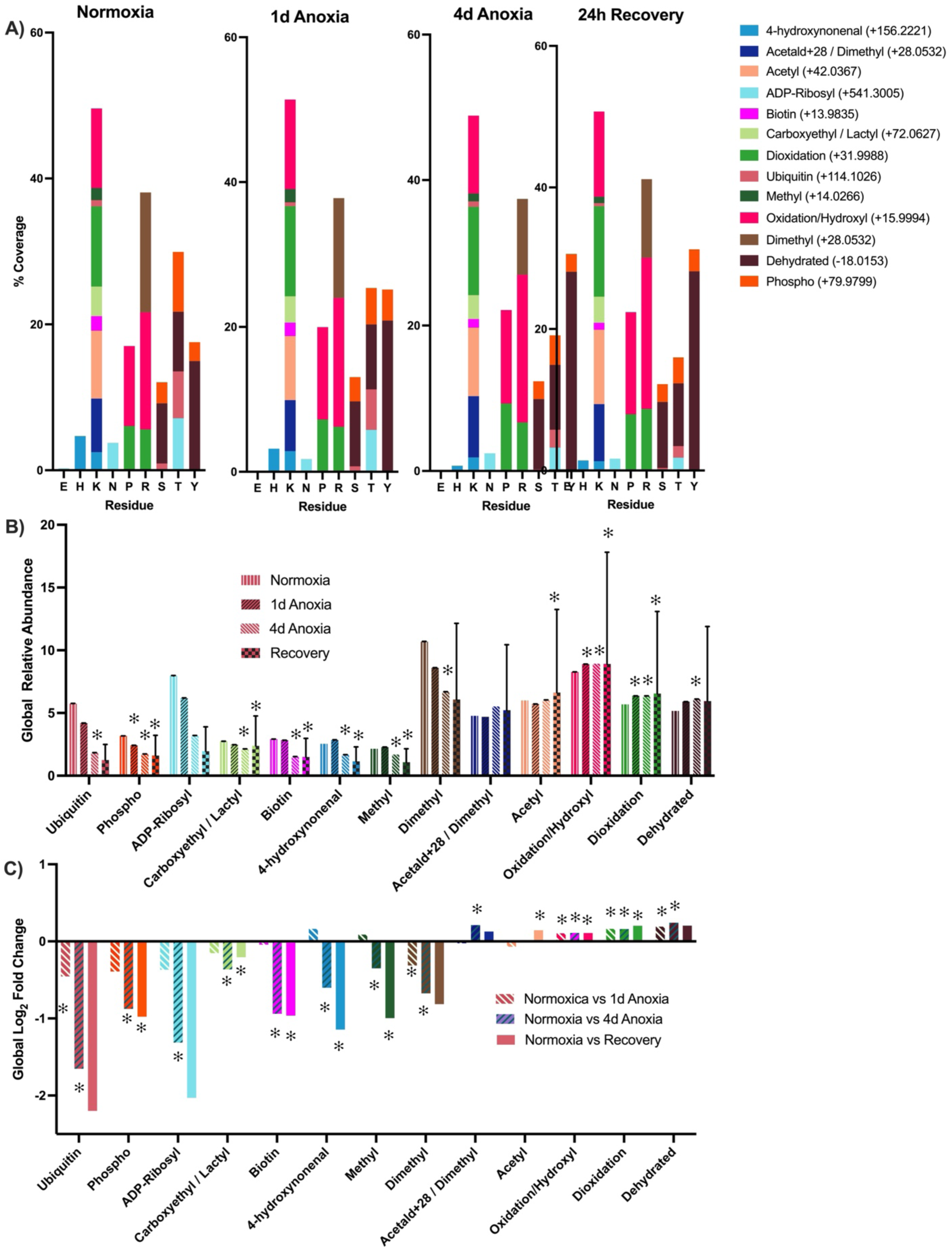
Global changes in the histone PTM landscape of WS40NE cells during anoxia and aerobic recovery from anoxia. **(A)** Thirteen types of hPTMs were detected across nine types of amino acid residues in WS40NE cells. The average mass as reported by Unimod is listed beside each PTM. While most residue types are unmodified, lysine (K) residues were the most consistently modified (54.7 - 58.3%) across the different experimental conditions. **(B)** An ANOVA with Dunnett’s multiple comparison test applied was used to determine significance of all conditions compared to normoxia (error bars represent standard deviation). From a global perspective, the abundance of most hPTMs decreases in response to anoxia and fails to return to normoxic values after 1 d of aerobic recovery. Acetylation, oxidation, dioxidation and dehydration increase modestly in response to anoxia. **(C)** Fold change data were calculated from the global mean abundance data for each hPTM. A two-tailed t-test was used to compare the relative abundance of hPTMs in all conditions with the Benjamini-Hochberg correction applied to account for multiple hypothesis testing.

### Characterization of histone H1

Six H1 proteins were identified: (**A**) H1-like (XP_013879563.1), (**B**) H1-like (XP_013855658.1), (**C**) H1-like (XP_013878240.1), (**D**) H1-like (XP_013888293.1), (**E**) H1-gamma late-like (XP_013861857.1), and (**F**) H1.0-B (XP_013863633.1). Thirteen types of PTMs were identified across these isoforms (Figs. 5A). Interestingly, H1-like (XP_013879563.1) appears to carry more modifications compared to the other isoforms even when percent coverage is taken into account (Figs. 1, 3D, 5C). Nine of the 15 modifications with an average relative abundance of exactly 100% in all samples are H1 isoforms, with H1-gamma late-like (XP_013861857.1) having 8 of the 10 (Table 1).

**Figure 5.**
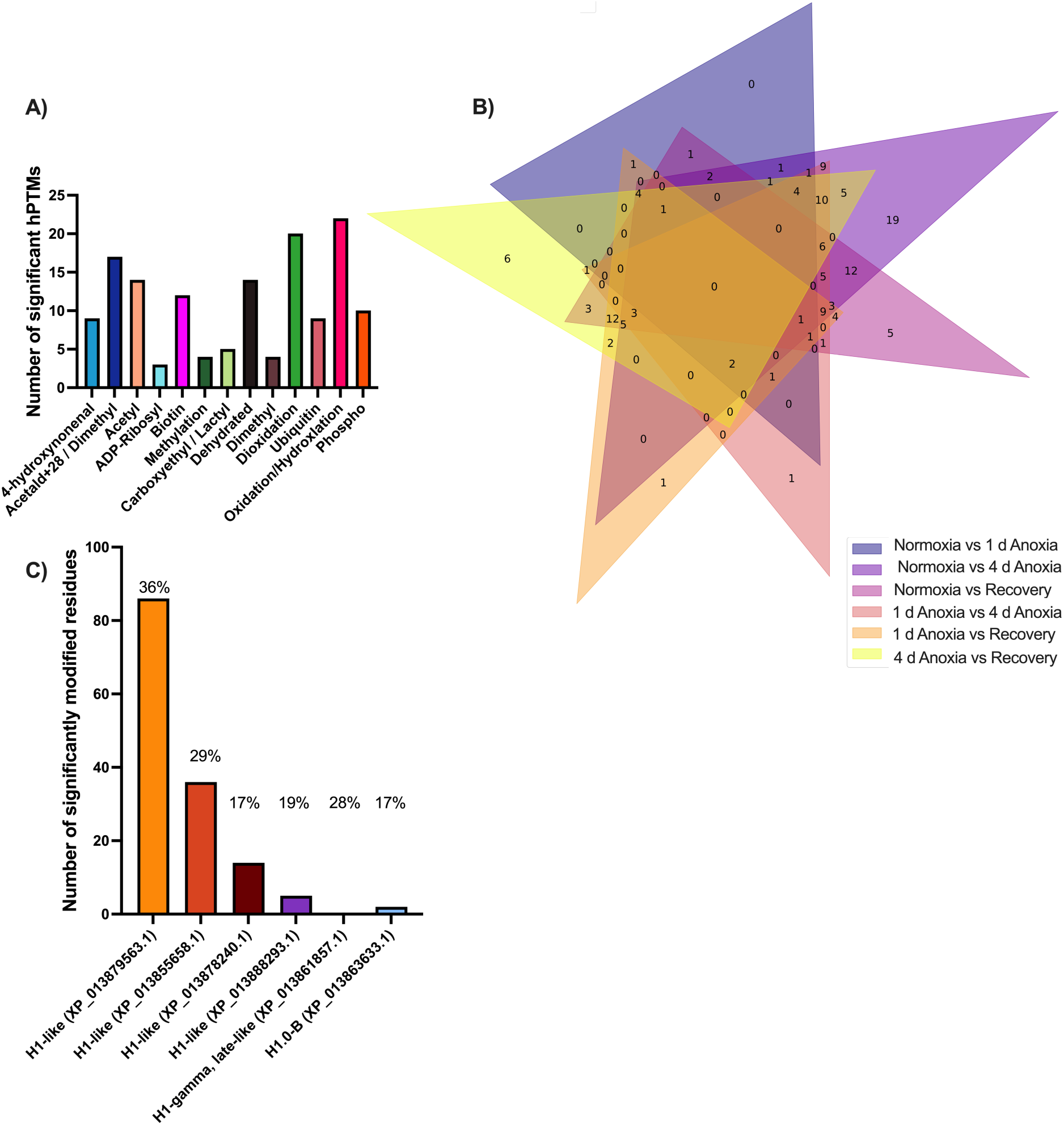
Significant differential histone post-translational modifications on histone H1 isoforms in WS40NE cells. (**A)** The most commonly significant differential hPTM across H1 isoforms were oxidation (22 hPTMs), dioxidation (20 hPTMs), and acetald+28/dimethylation (17 hPTMs). (**B**) Venn diagram illustrating the occurrence of unique and biologically-relevant hPTMs on H1 histone isoforms across experimental treatment groups. The Venn diagram is schematic and non-proportional. The numbers within each colored section indicate the number of significantly differentially expressed histone modifications shared among the indicated pairwise comparisons. (**C**) Five of the six H1 histone isoforms had residues with significantly different hPTM modifications in response to anoxia. The percentage above each bar represents the peptide sequence coverage of that isoform. Note that H1-gamma late-like is modified, but those modifications are stable across experimental treatment groups.

All six H1 isoforms had residues with significant hPTM modifications, while 5 isoforms had at least one hPTM that changed in relative abundance in response to experimental treatments. One hundred and forty-three significant hPTMs were present on H1 isoforms in at least one condition (Fig. 5). The most common differential modifications in response to the experimental treatments across H1 isoforms were oxidation (22 significant unique hPTMs), dioxidation (20 significant unique hPTMs), and acetald+28/dimethylation (17 significant unique hPTMs) (Fig. 5). Six highly condition-dependent hPTMs were significant across five different comparisons: H1-like (XP_013879563.1) 17K acetylation, 62R oxidation/hydroxylation, and 63S phosphorylation; H1-like (XP_013855658.1) 51K 4-hydroxynonenal; and H1.0-B (XP_013863633.1) 19K 4-hydroxynonenal and 25K acetylation (Fig. 6 and Table 2).

**Figure 6.**
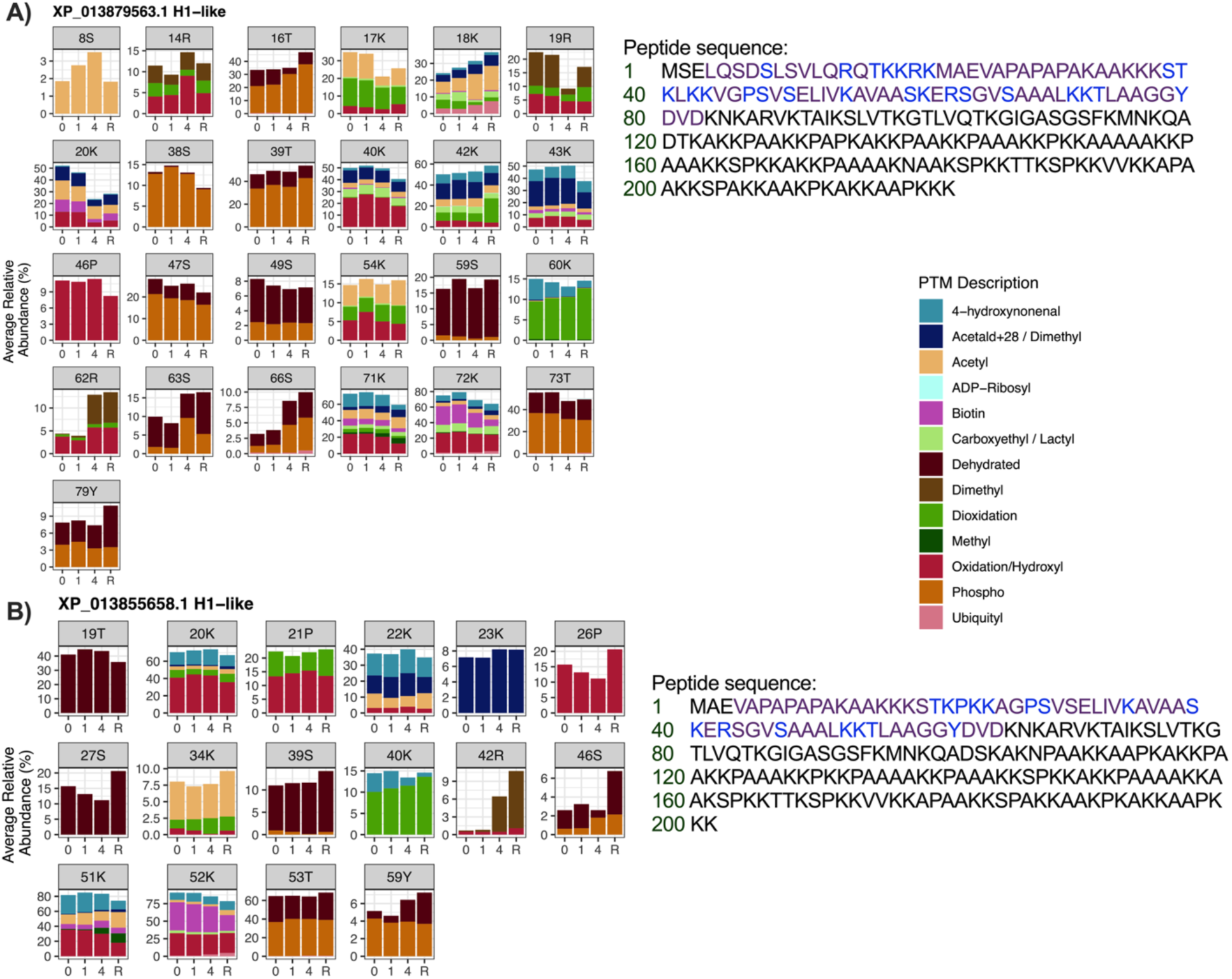
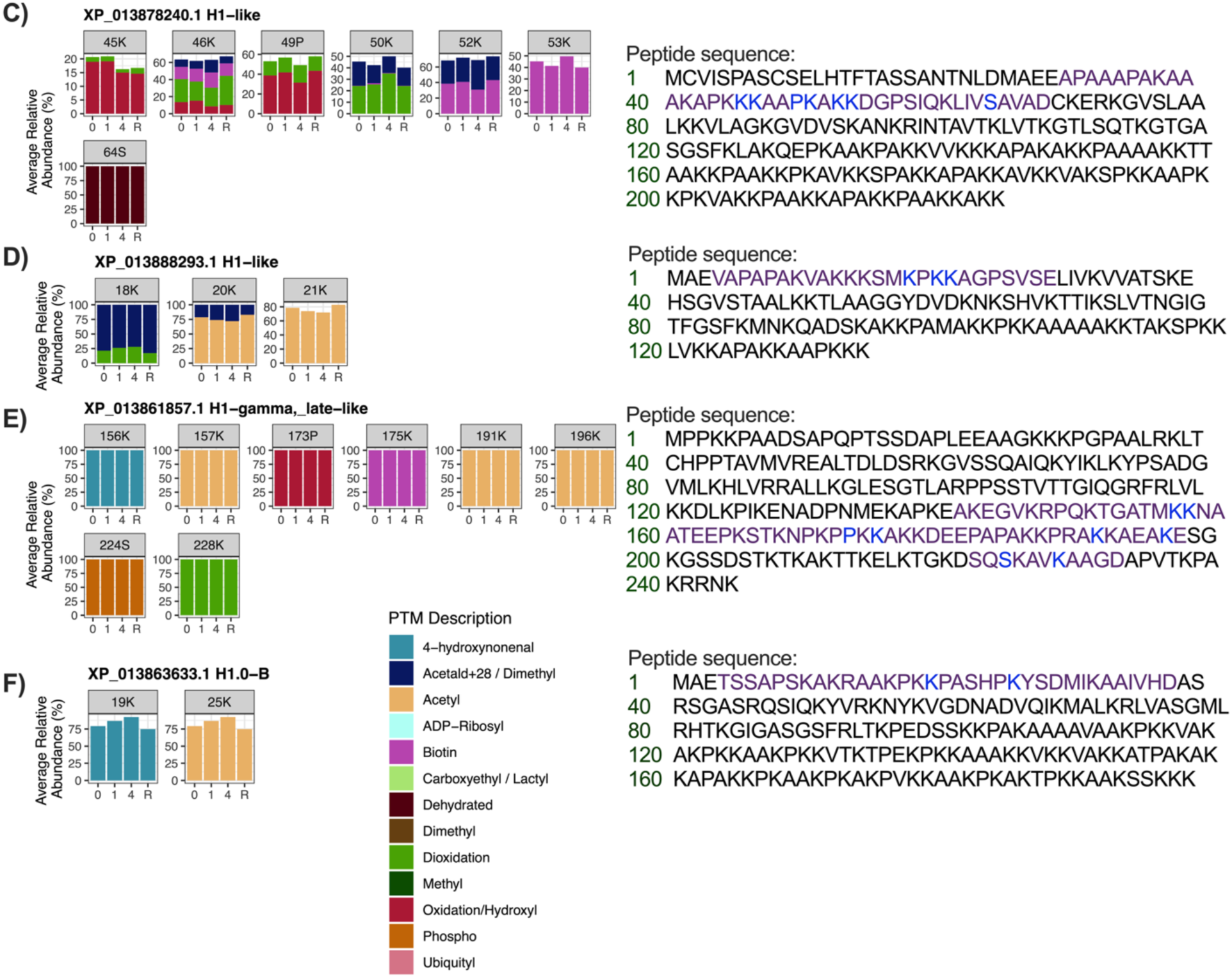
Characterization of post-translational modifications on Histone H1 isoforms in WS40NE cells exposed to anoxia and aerobic recovery from anoxia. Isoforms represented are (**A**) H1-like (XP_013879563.1), (**B**) H1-like (XP_013855658.1), (**C**) H1-like (XP_013878240.1), (**D**) H1-like (XP_013888293.1), (**E**) H1-gamma late-like (XP_013861857.1), and (**F**) H1.0-B (XP_013863633.1). Each panel represents a different amino acid position where at least one hPTM was detected. Amino acid residues are abbreviated using their one letter code. The x-axis displays the condition of the samples: 0 d in anoxia (0), 1 d in anoxia (1), 4 d in anoxia (4), or 1 d of aerobic recovery from 4 d of anoxia (R). The y-axis scales differ across panels to better visualize changes in low abundance hPTMs. Detected (purple) and modified (blue) peptides and the entire peptide sequences for each isoform are listed beside the panels. On each row, the number of the corresponding residue is listed in green.

### Characterization of histone H2A

Nine H2A proteins were identified: (**A**) H2A (XP_013874651.1), (**B**) H2A-like (XP_013879560.1), (**C**) H2A-like (XP_013878193.1), (**D**) macro-H2A.2 (XP_013858189.1), (**E**) core histone macro-H2A.1 (XP_013877703.1), (**F**) H2A.V (XP_013854970.1), (**G**) H2A.Z (XP_013858796.1), (**H**) H2AX-like (XP_013887205.1), and (**I**) H2AX-like (XP_013862107.1/ XP_013878242.1). Thirteen types of PTMs were identified across these isoforms (Fig. 7A) with 257 hPTMs significantly differentially abundant on H2A isoforms in at least one condition (Fig. 7B). All nine H2A isoforms had residues with significant hPTM modifications, with H2A (XP_013874651.1) and H2A-like (XP_013879560.1) appearing to have a high proportion of hPTMs for their percent coverage in the data (Fig. 7C). The top three most common hPTMs that were significantly different in abundance in response to experimental treatments across H2A isoforms were dioxidation (45 significant unique hPTMs), oxidation (38 significant unique hPTMs), and dehydration (25 significant unique hPTMs) (Fig. 7A). Two highly condition-dependent hPTMs were significantly different across six different comparisons: H2A-like (XP_013879560.1) 123K carboxyethylation/lactylation (Figs. 7E, 8B) and H2A-like (XP_013878193.1) 102S dehydration (Figs. 7D, 8C). Two additional highly condition-dependent hPTMs were significantly different across five different comparisons: H2A.V (XP_013854970.1)121K biotinylation and 122K dioxidation (Fig. 9A, Table 2).

**Figure 7.**
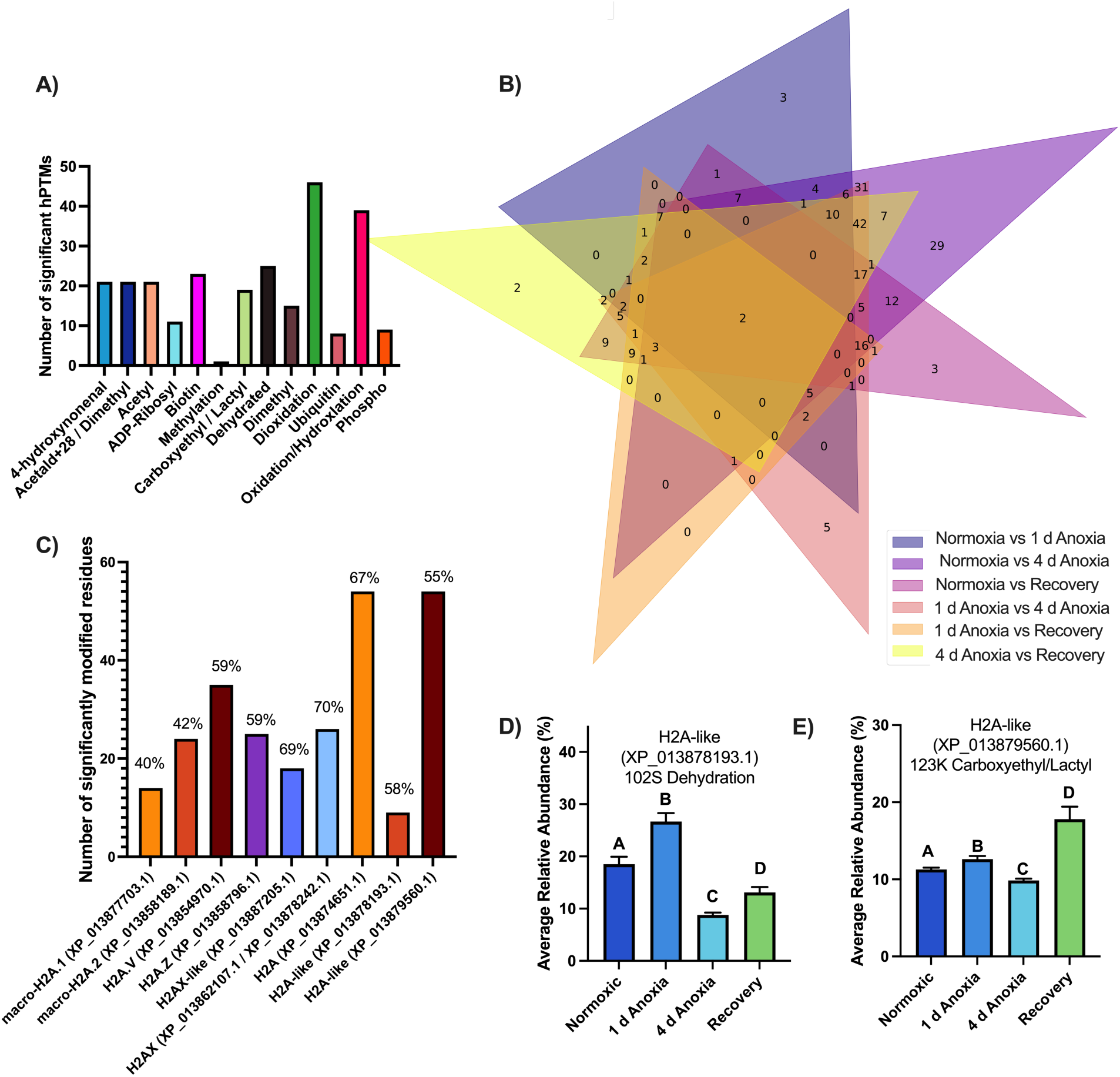
Significant differential post-translational modifications on histone H2A isoforms in WS40NE cells exposed to anoxia and aerobic recovery from anoxia. **(A)** The top three most commonly significant differential hPTMs across histone H2A isoforms were dioxidation (45 hPTMs), oxidation (38 hPTMs), and dehydration (25 hPTMs). (**B**) Venn diagram illustrating the occurrence of unique and biologically-relevant hPTMs on H2A histone isoforms across experimental treatment groups. The Venn diagram is schematic and non-proportional. The numbers within each colored section indicate the number of significantly differentially expressed histone modifications shared among the indicated pairwise comparisons. (**C**) At least nine H2A isoforms had residues that were significantly modified. The percentage above each bar represents the peptide sequence coverage of that isoform. Note that two isoforms appear to be highly modified based on their percent sequence coverage. Two specific hPTMs were significant across all six pairwise comparisons: **(D)** H2A-like (XP_013878193.1) 102S dehydration and **(E)** H2A-like (XP_013879560.1) 123K carboxyethylation/lactylation. Bars represent means ± SEM (n=11-12).

**Figure 8.**
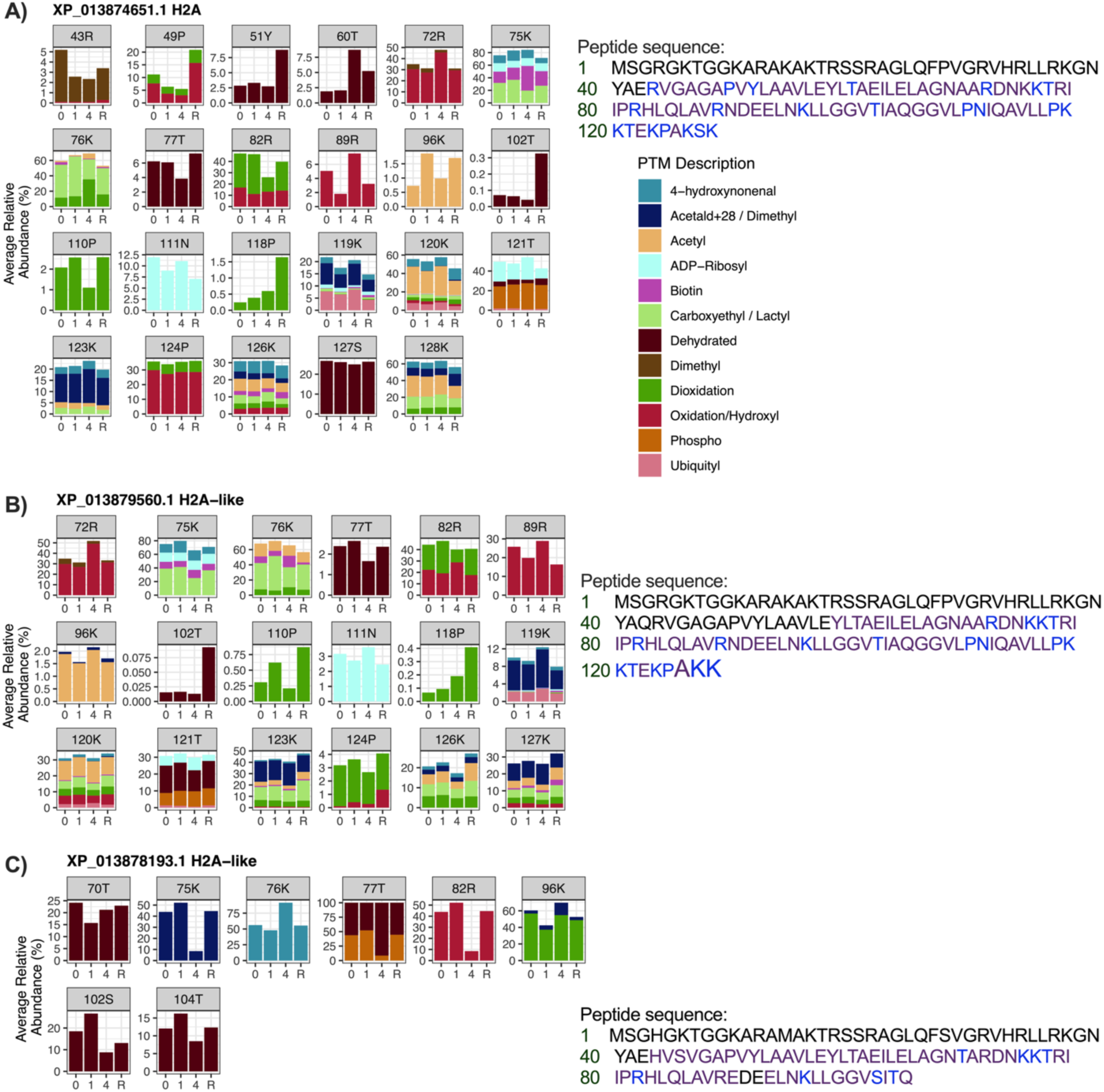
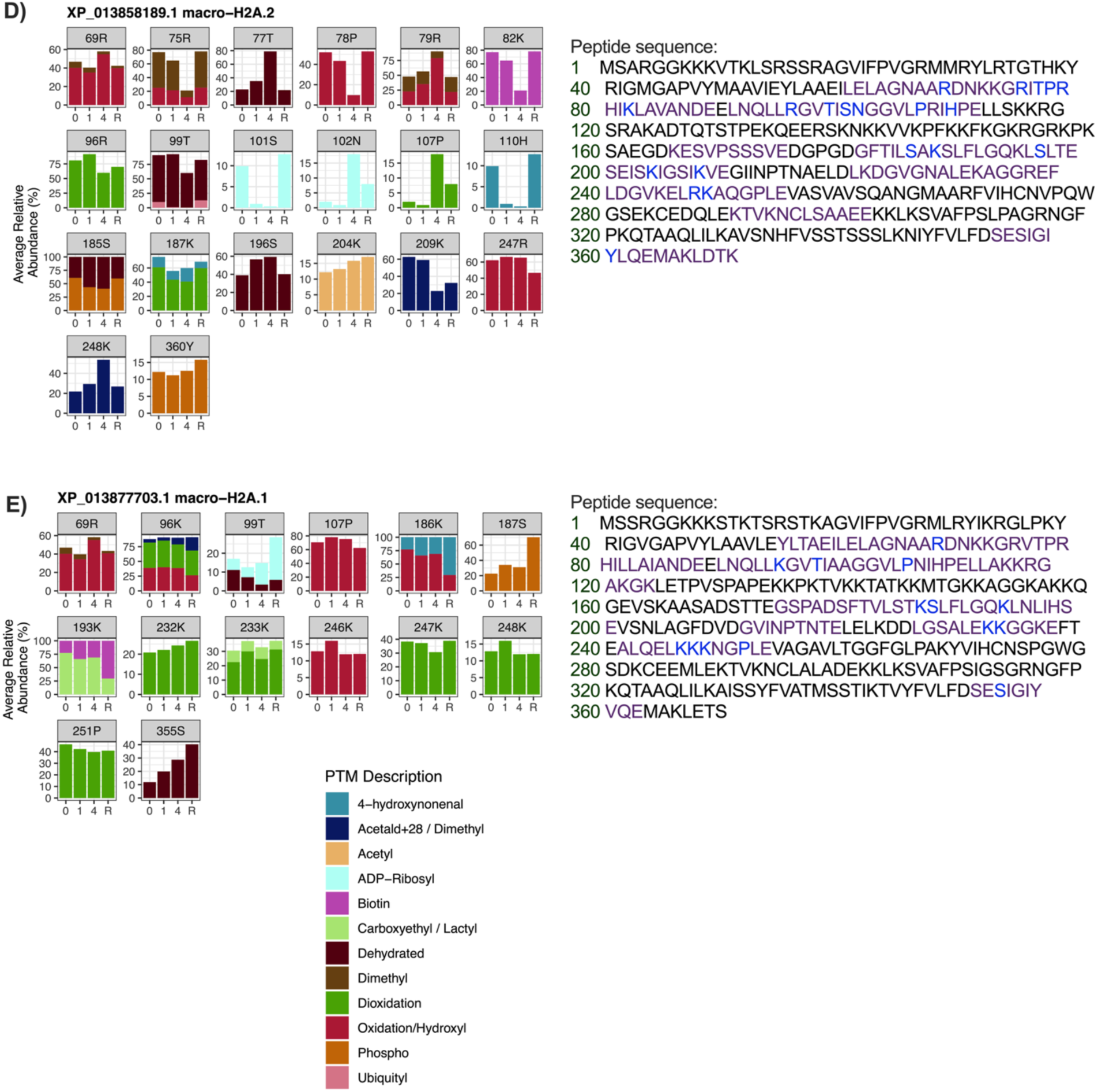
Characterization of the post-translational modifications on histone H2A isoforms in WS40NE cells exposed to anoxia and aerobic recovery from anoxia. Isoforms represented are (**A**) H2A (XP_013874651.1), (**B**) H2A-like (XP_013879560.1), (**C**) H2A-like (XP_013878193.1), (**D**) macro-H2A.2 (XP_013858189.1), and (**E**) macro-H2A.1 (XP_0138777031.1). Each panel represents a different amino acid position where at least one hPTM was detected. Amino acid residues are abbreviated using their one letter code. The x-axis displays the condition of the samples: 0 d in anoxia (0), 1 d in anoxia (1), 4 d in anoxia (4), or 1 d of aerobic recovery after 4 days in anoxia (R). The y-axis scales differ across panels to better visualize changes in low abundance hPTMs. Detected (purple) and modified (blue) peptides and the entire peptide sequences are listed beside the panels. On each row, the number of the corresponding residue is listed in green.

**Figure 9.**
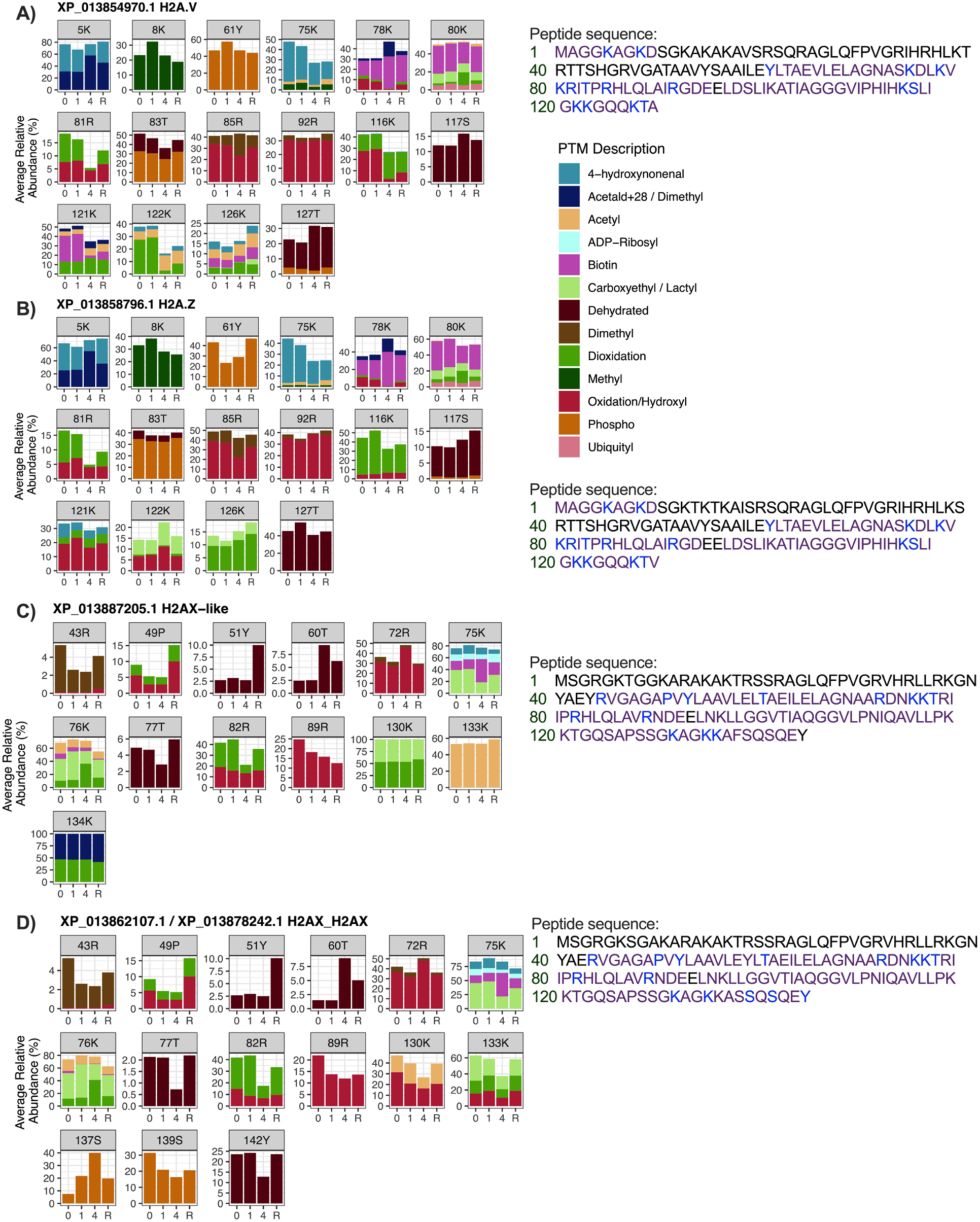
Characterization of the post-translational modifications on histone H2A isoforms in WS40NE cells exposed to anoxia and aerobic recovery from anoxia. (**A**) H2A.V (XP_013854970.1), (**B**) H2A.Z (XP_013858796.1), (**C**) H2AX-like (XP_013887205.1), and (**D**) H2AX-like (XP_013862107.1/ XP_013878242.1). Each panel represents a different amino acid position where at least one hPTM was detected. Amino acid residues are abbreviated using their one letter code. The x-axis displays the condition of the samples: 0 d in anoxia (0), 1 d in anoxia (1), 4 d in anoxia (4), or 1 d of aerobic recovery following 4 d of anoxia (R). The y-axis scales differ across panels to better visualize changes in low abundance hPTMs. Detected (purple) and modified (blue) peptides and the entire peptide sequences are listed beside the panels. On each row, the number of the corresponding residue is listed in green.

### Characterization of histone H2B

At least five H2B proteins were identified: (**A**) H2B_1/2-like (XP_013874662.1), (**B**) H2B_1/2 (XP_013879561.1 / XP_013878239.1 / XP_013878163.1 / XP_013869445.1), (**C**) H2B_1/2-like (XP_013857090.1), (**D**) H2B.L4-like (XP_013885354.1), and (**E**) H2B.3-like (XP_013886452.1). Thirteen types of PTMs were identified across these isoforms (Fig. 10A). Three hundred and twenty-seven significant hPTMs were present across H2B isoforms in at least one condition, making this the most significantly modified class of histones in this study (Fig. 10B). All identified H2B isoforms had at least one residue with a significant hPTM, while 4 of the 5 were highly modified (Fig. 10C). The top three most commonly significant differential hPTMs across H2B isoforms were phosphorylation (58 significant unique hPTMs), dehydration (49 significant unique hPTMs), and oxidation/hydroxylation (35 significant unique hPTMs) (Fig. 10A).The following hPTMs displayed an average relative abundance of exactly 100% in all samples: H2B.L4-like (XP_013885354.1) 31Y, 43T, 46S, and 55S dehydration (Fig. 11, Table 1).

**Figure 10.**
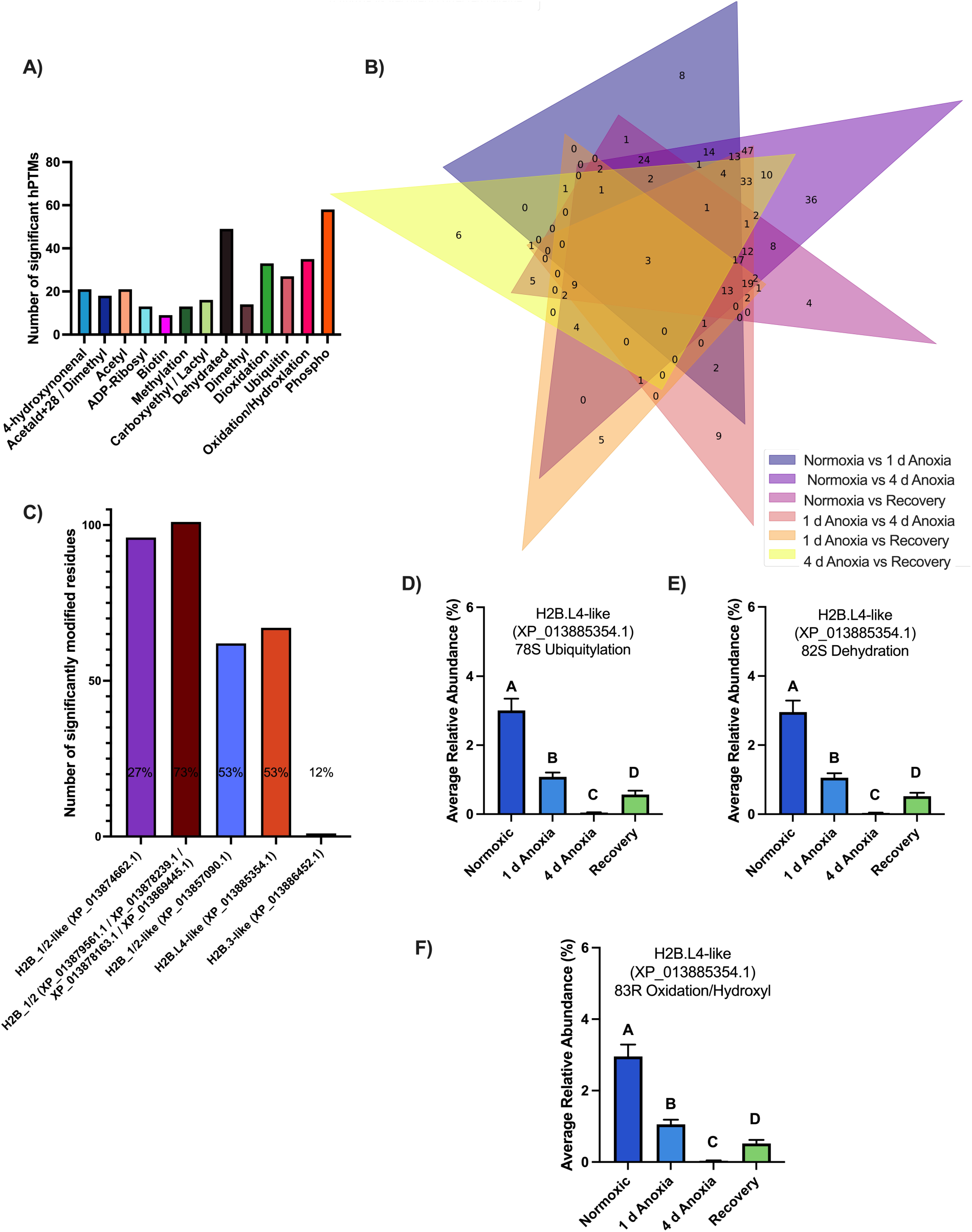
Significant differential post-translational modifications on histone H2B isoforms in WS40NE cells exposed to anoxia and aerobic recovery from anoxia. **(A)** The top three most commonly significant differential hPTMs across H2B isoforms were phosphorylation (58 hPTMs), dehydration (48 hPTMs), and oxidation/hydroxylation (35 hPTMs). (**B**) Venn diagram illustrating the occurrence of unique and biologically-relevant hPTMs on H2B histone isoforms across experimental treatment groups. The Venn diagram is schematic and non-proportional. The numbers within each colored section indicate the number of significantly differentially expressed histone modifications shared among the indicated pairwise comparisons. (**C**) At least five H2B isoforms had residues with significant hPTMs. The percentage above each bar represents the peptide sequence coverage of that isoform. Three hPTMs on H2B.L4-like (XP_013885354.1) were highly condition dependent across all six comparisons: **(D)** 78S ubiquitylation, **(E)** 82S dehydration, and **(F)** 83R oxidation/hydroxylation. Bars represent the mean ± SEM (n=11-12).

**Figure 11.**
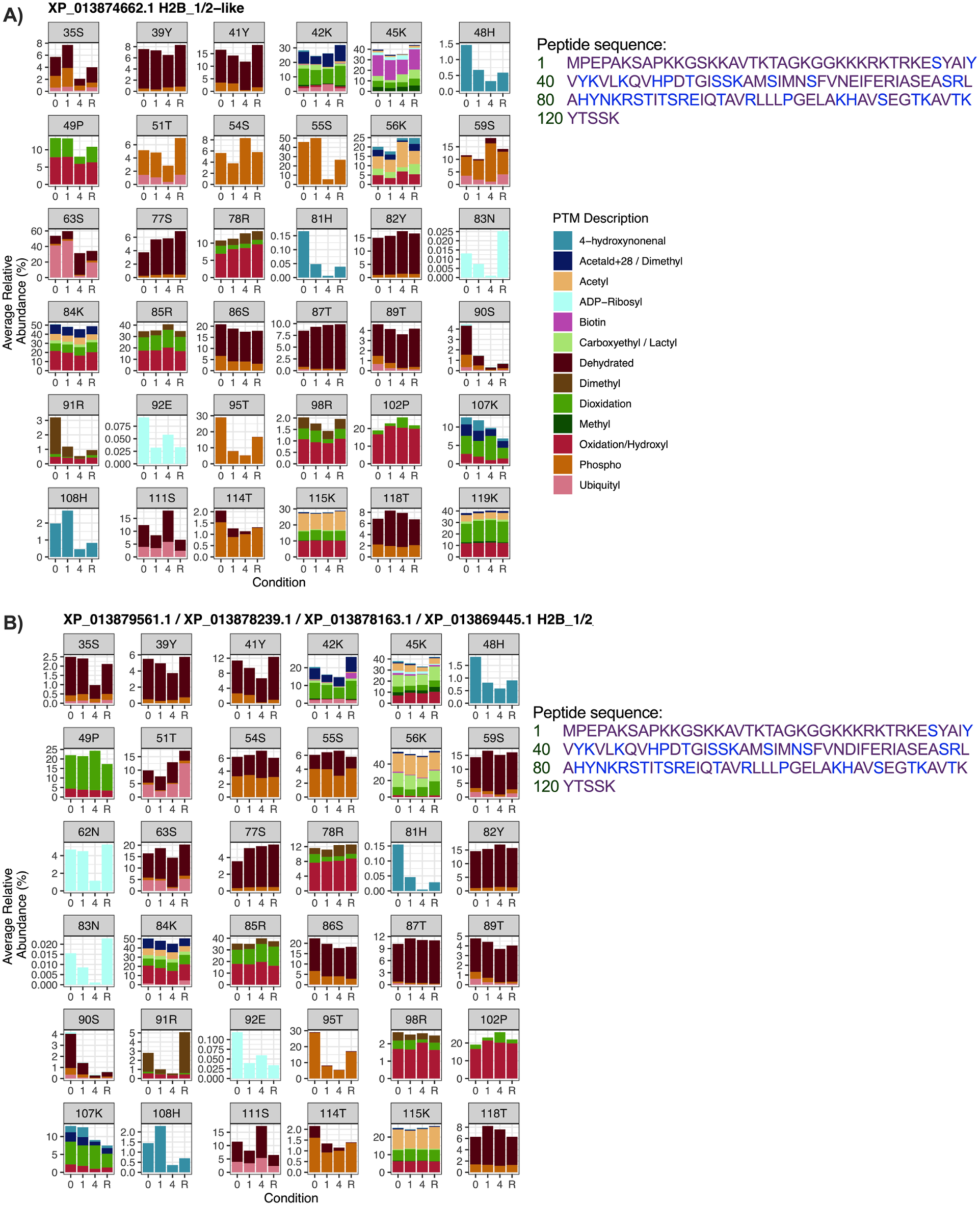
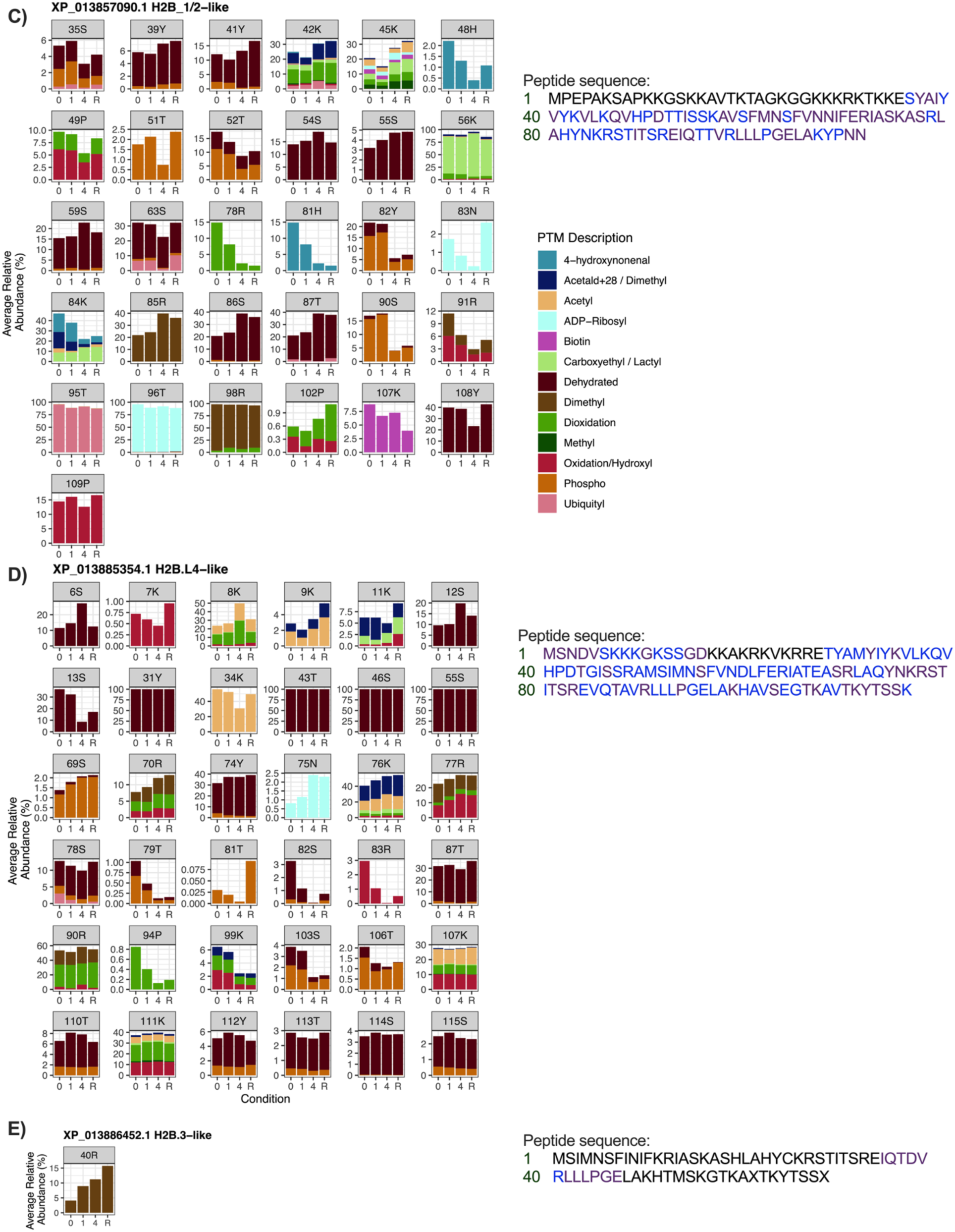
Characterization of the post-translational modifications on histone H2B isoforms in WS40NE cells exposed to anoxia and aerobic recovery from anoxia. Isoforms represented are (**A**) H2B_1/2-like (XP_013874662.1), (**B**) H2B_1/2 (XP_013879561.1/ XP_013878239.1 / XP_013878163.1 / XP_013869445.1), (**C**) H2B_1/2-like (XP_013857090.1), (**D**) H2B.L4-like (XP_013885354.1) and (**E**) H2B.3-like (XP_013886452.1). Each panel represents a different amino acid position where at least one hPTM was detected. Amino acid residues are abbreviated using their one letter code. The x-axis displays the condition of the samples: 0 d in anoxia (0), 1 d in anoxia (1), 4 d in anoxia (4), or 1 d of aerobic recovery after 4 d of anoxia (R). The y-axis scales differ across panels to better visualize changes in low abundance hPTMs. Detected (purple) and modified (blue) peptides and the entire peptide sequences are listed beside the panels. On each row, the number of the corresponding residue is listed in green.

Three hPTMs on H2B.L4-like (XP_013885354.1) were highly condition dependent across all six comparisons: 78S ubiquitylation, 82S dehydration, and 83R oxidation/hydroxylation (Fig. 10D,E,F). An additional twenty-three hPTMs were highly condition dependent across five different comparisons on four different histone isoforms: eight on H2B_1/2-like (XP_013874662.1), seven on H2B_1/2 (XP_013879561.1 / XP_013878239.1 / XP_013878163.1 / XP_013869445.1), four on H2B_1/2-like (XP_013857090.1), and four on H2B.L4-like (XP_013885354.1) (Fig. 11, Table 2). Two of these highly condition-dependent modifications on H2B.L4-like (XP_013885354.1) occur on the same residue: 79T dehydration and phosphorylation (Fig. 11D, Table 2).

### Characterization of histone H3

At least three H3 proteins were identified: (**A**) H3.3 (XP_013887037.1 / XP_013885059.1 / XP_013879174.1 / XP_013870443.1), (**B**) H3-like (XP_013879564.1), and (**C**) H3-like (XP_013878241.1). All identified H3 isoforms had at least one residue with a significant hPTM, with 12 types of hPTMs identified across these isoforms (Fig. 12A). The most common hPTMs that differ significantly across experimental treatments on H3 isoforms were oxidation/hydroxylation (23 significant unique hPTMs) and dioxidation (18 significant unique hPTMs) (Fig. 12A). Eighty-nine hPTMs were present across all three H3 isoforms in at least one condition (Fig. 12B). Two hPTMs displayed an average relative abundance of exactly 100% in all samples: H3-like (XP_013878241.1) 17K and 24R oxidation/hydroxylation (Fig. 13C, Table 1).

**Figure 12.**
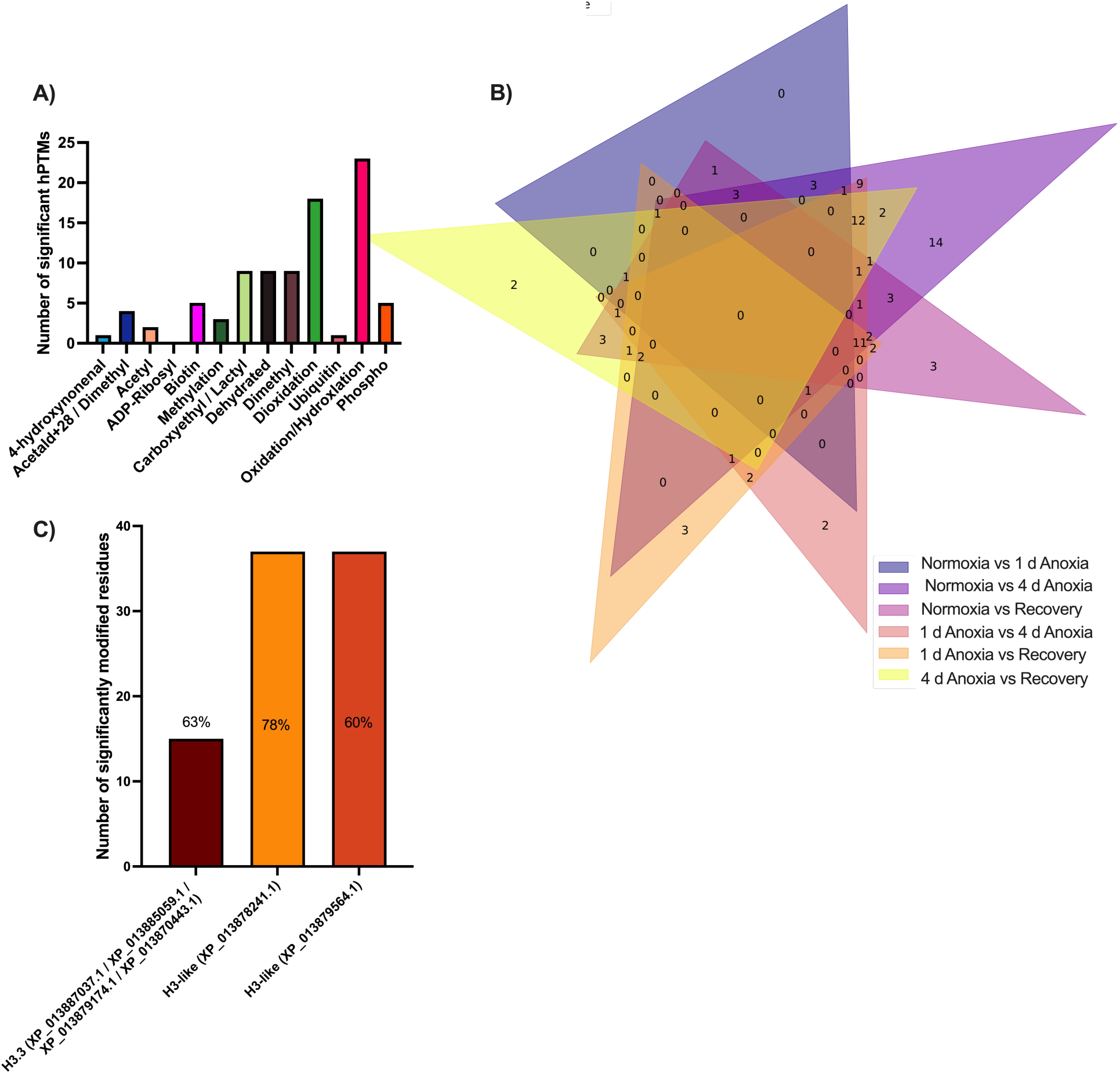
Significant differential post-translational modifications on histone H3 isoforms in WS40NE cells exposed to anoxia and aerobic recovery from anoxia. **(A)** The most commonly significant differential hPTMs across H3 isoforms were oxidation (23 hPTMs) and dioxidation (18 hPTMs). (**B**) Venn diagram illustrating the occurrence of unique and biologically-relevant hPTMs on H3 histone isoforms across experimental treatment groups. The Venn diagram is schematic and non-proportional. The numbers within each colored section indicate the number of significantly differentially expressed histone modifications shared among the indicated pairwise comparisons. (**C**) All identified H3 isoforms had residues with significant hPTM modifications. The percentage above each bar represents the peptide sequence coverage of that isoform (n=11-12).

**Figure 13.**
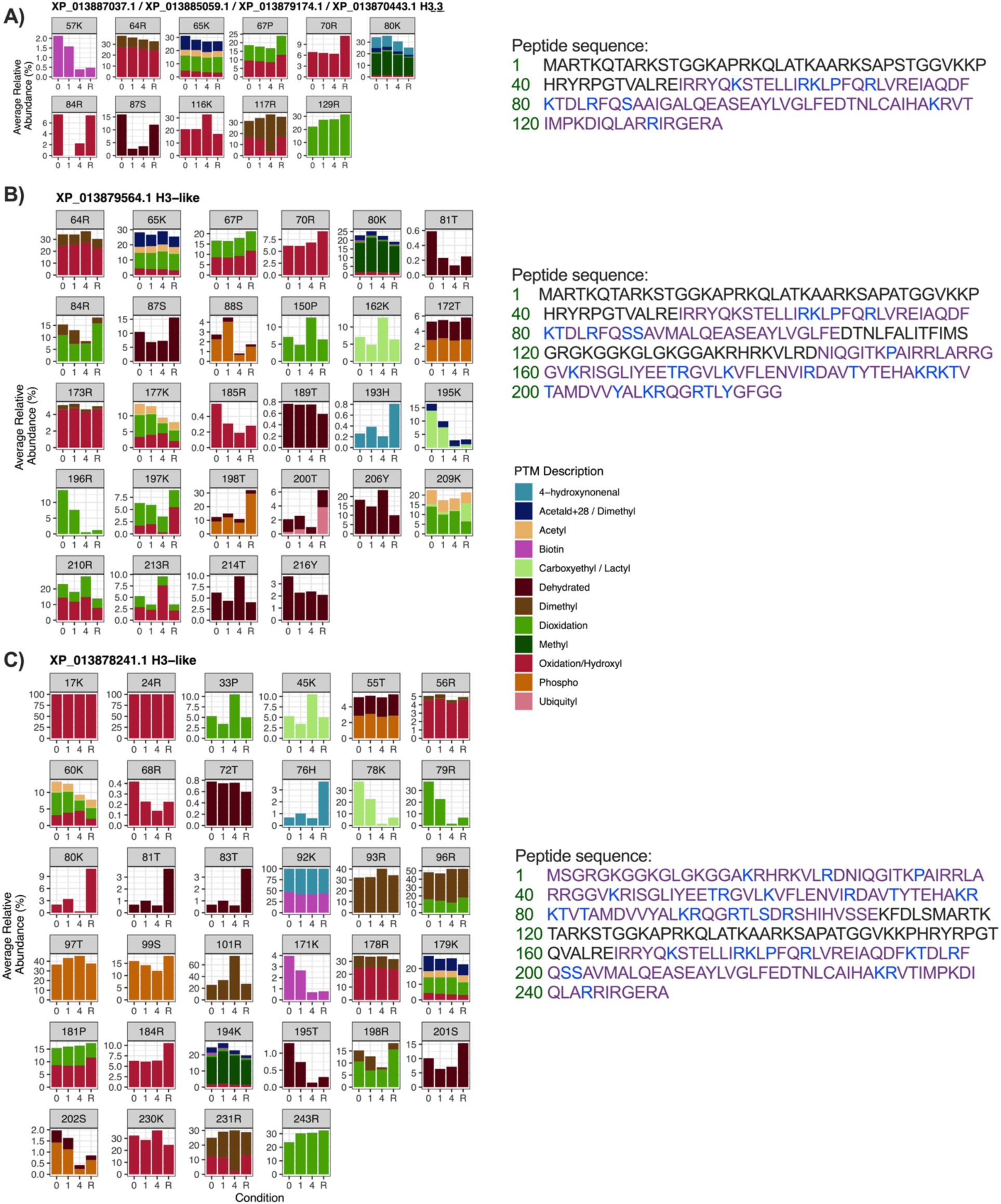
Characterization of the post-translational modifications on histone H3 isoforms in WS40NE cells exposed to anoxia and aerobic recovery from anoxia. Isoforms represented are (**A**) H3.3 (XP_013887037.1/ XP_013885059.1/ XP_013879174.1 / XP_013870443.1), (**B**) H3-like (XP_013879564.1), and (**C**) H3-like (XP_013878241.1). Each panel represents a different amino acid position where at least one hPTM was detected. Amino acid residues are abbreviated using their one letter code. The x-axis displays the condition of the samples: 0 d in anoxia (0), 1 d in anoxia (1), 4 d in anoxia (4), or 1 d aerobic recovery after 4 d of anoxia (R). The y-axis scales differ across panels to better visualize changes in low abundance hPTMs. Detected (purple) and modified (blue) peptides and the entire peptide sequences are listed beside the panels. On each row, the number of the corresponding residue is listed in green.

### Extracellular Lactate Accumulation

While previous studies have explored accumulation of lactate during exposure to anoxia in WS40NE cells, this was the first study to monitor how this lactate accumulation persisted during recovery (Fig. 14). As previously shown, anoxic WS40NE cells accumulate extracellular lactate at a significantly higher rate than normoxic cells. After 1 d of aerobic recovery, this trend persists.

**Figure 14.**
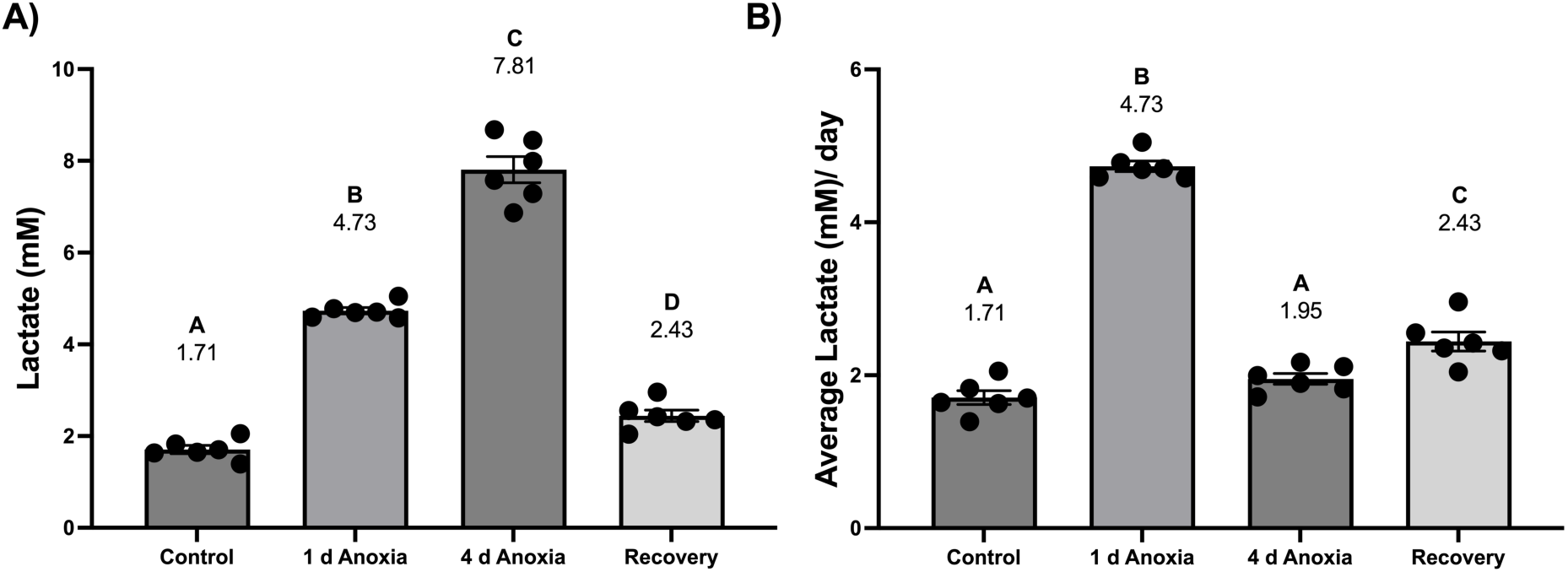
Extracellular lactate accumulation in WS40NE cells exposed to anoxia and recovery from anoxia. (**A**) Total media lactate concentration (mM) during exposure to anoxia and recovery from anoxia. Due to the recovery media change, the data represents only 1 day of lactate accumulation in normoxic recovery conditions. (**B**) The average rate of lactate accumulation (mM/day) in the same samples. Different letters indicate significant differences in means within each graph (one-way ANOVA with Tukey’s HSD test p > 0.05, n = 6).

## Discussion

Histone isoforms and post-translational modifications change in response to anoxia and aerobic recovery from anoxia in WS40NE cells. Of the 1043 unique biologically relevant hPTMs detected, 816 hPTMs were significantly modified in at least one condition. All identifiable histone groups showed significant post-translational modifications in response to changing oxygen availability. H2B was the most significantly modified histone group with 327 significant differential hPTMs present in at least one condition. H2B_1/2 (XP_013879561.1 / XP_013878239.1 / XP_013878163.1 / XP_013869445.1) was the most significantly modified histone, with 101 significant modifications. Interestingly, highly condition-dependent hPTMs often occurred on adjacent residues, with two highly condition-dependent hPTMs sharing a single residue. This suggests that histones of WS40NE cells have highly modifiable peptide regions that are responsive to changes in oxygen availability and likely help to coordinate changes in gene expression associated with their extreme anoxia tolerance. In many aspects, this response is consistent with hypoxic responses in hPTMs observed in human cells [39, 40], including several stress related hPTMs, such as oxidation/hydroxylation and dioxidation, increasing globally in response to anoxic stress and aerobic recovery due to increased rates of reactive oxygen species (ROS) production [41, 42]. However, WS40NE cells demonstrate several unexpected responses to changing aerobic conditions, including changes in histone isoform abundance and several unique trends in hPTMs.

### Limitations and precautions in the interpretation of these data

One limitation of this study is the lack of detection of histone H4. This result is consistent with previous studies using these methods in *A. limnaeus* [34] and in other fish species [20, 43]. As previously discussed, this is likely due to the specific nature of the single histone H4 sequence in *A. limnaeus* and the limited number of potential ions available for detection using these methodologies. In addition, it is important to note that while there is a great deal of sequence conservation for histone across species, some sequences in *A. limnaeus* contain long N-terminal tails that make direct comparisons of the “classic” histone modifications difficult (see discussion in [34]). It also means that for those isoforms, what is considered a “tail” in a mammalian protein becomes more central in *A. limnaeus*, despite the tail portion of the protein being highly modified (e.g. Figure 13c). The significance of these differences is unknown but should be considered when comparing these data to those in other species.

### Changes in histone isoform abundance

Several histone isoforms change in abundance in response to anoxia. While, to our knowledge, no study has identified anoxia-dependent changes in histone isoform abundance, there are several studies that document changes in histone isoforms in response to heat stress, hypoxia, ischemia, and oxidative stress [40, 44–46]. Notably, one of the histone variants to show significant changes in abundance in our study H3.3, is generally deposited via histone cell cycle regulator (HIRA) in a replication-independent manner [40, 47], suggesting that its hPTM and isoform abundance changes are not due to the cell division that can occur during anoxia [37]. The increase in histone H3.3 observed in this study is consistent with a role for “priming” the cells for a robust gene expression response to stress as has been shown in human lung fibroblasts [40]. Importantly, histone H3.3 expression responds within 1 day of anoxia and does not immediately return to pre-anoxic levels after 1 day of recovery, suggesting a prolonged change in poise of the chromatin. H3.3 is typically associated with euchromatin and active regulatory elements [48], suggesting that the genomic location of the histone H3.3 isoforms may indicate genes that are immediately responsible for supporting cellular functions in the absence of oxygen, or may represent genes primed and ready to be expressed during recovery from anoxia. If future studies can identify the genomic location of the H3.3 isoforms, these genes would be prime candidates for explaining anoxic-preconditioning in embryos of *A. limnaeus* [49].

The other differentially abundant isoforms are most closely related to canonical H2A and H2B isoforms [50], which can be exchanged via replication dependent mechanisms [51]. However, these isoforms can also be exchanged via replication independent mechanisms including complexes formed by histone chaperone nucleosome assembly protein 1 (NAP-1) or facilitates chromatin transcription (FACT) protein [52–54]. Given that FACT can be recruited during hypoxia to modulate both histone ubiquitylation and transcriptional activation of hypoxia-induced genes, the ability of these complexes to exchange histone isoforms could very well also be a part of the anoxic stress response [55]. H2A/H2B dimers flank entry and exit portals for DNA in the nucleosome, and thus changes to these histone isoforms are highly likely to impact gene expression [51, 56]. In WS40NE cells, one histone H2A isoform decreases within 1 day of anoxia, while two histone H2B isoforms decrease, but only after 4 days of anoxia, suggesting unique regulation and usage of these isoforms in response to anoxia. Histone H2A isoforms have been shown to increase after exposure to hypoxia and ischemia in some mammalian models [57, 58], a result that is opposite of that in WS40NE cells. The reason for this shift and their potential for altering gene expression is worthy of future studies.

### Global decreases in multiple hPTMs deviate from expected patterns

Global changes to hPTMs have been recorded in mammalian cells in response to hypoxia; large changes in modifications of H3 and H4 histones are seen in HEK 293T, HeLa and RKO cells with smaller changes occurring on H2A and H2B histones [39]. However, few studies have investigated anoxia-induced changes in hPTMs in a modification-specific way [34]. Interestingly, global methylation in WS40NE cells is stable for the first day of anoxia, but then significantly decreases during 4 d anoxia and remains low for 1 d of aerobic recovery. As H3K4me3 and H3K36me3 increase within 1 h during hypoxia in mammalian systems, we expected global methylation in *A. limnaeus* to reflect similar changes [28]. Increases in methylation in hypoxic mammalian systems is thought to be due to decreased activity of histone demethylases when oxygen is limiting, mainly because many of these enzymes use molecular oxygen as a substrate, and at least one histone demethylase appears to be upregulated in a hypoxia-inducible factor dependent manner, presumably to combat the overall loss of demethylase activity during hypoxia [59]. Thus, the large and sustained decrease in global methylation exhibited in killifish WS40NE cells is an unexpected result when referenced to the mammalian literature. Histone methylation, including di- and trimethylation, is associated with transcriptional activation and repression in a residue specific manner [60, 61], so it is difficult at this time to draw any conclusions for the role that decreased methylation might play. Of note, only one site of methylation in our dataset was highly condition dependent (H2B_1/2, XP_013874662.1 and XP_013869445.1, 124K). This residue does not correspond with any known mammalian hPTMs but may serve a similar function and should be investigated further.

Given its role in the regulation of DNA repair [62], it is surprising that global levels of ADP-ribosylation consistently decrease in anoxia and remain depressed during aerobic recovery. Most residue-specific patterns match this global pattern with either stable or decreasing levels of ADP-ribosylation during anoxia. However, in the cases where an increase occurs on a specific residue, it is almost always during aerobic recovery. This suggests a conserved role for ADP-ribosylation of histones associated with oxidative stress and the DNA damage response [62, 63]. However, PARPs including diphtheria-toxin-like ADP-ribose transferases (ARTDs) and cholera-toxin-like ARTCs can modify a variety of residues: lysine, aspartate, glutamate, serine, arginine, tyrosine, and histidine [64, 65]. Additionally, ADP-ribosylation can occur on arginine, serine, threonine, and cysteine residues via the action of sirtuins (SIRT4, SIRT6, and SIRT7) that perform mono (ADP-ribose) (MAR) addition [64]. Histone ADP-ribosylation is associated with a variety of functions beyond DNA damage response, including chromatin remodeling and facilitation of gene transcription by making chromatin more accessible [66, 67]. A decrease in ADP-ribosylation may be associated with a decrease in chromatin availability, decreasing the accessibility of the genome to polymerases and transcription factors. Thus, the overall decrease in histone ADP-ribosylation may support the downregulation of transcription associated with entrance into anoxia-induced quiescence.

Recent studies suggest lactate accumulation in mammalian cells may play a role in cell signaling and regulation through histone lactylation [68], including stimulating gene transcription in macrophages [69]. WS40NE cells accumulate extracellular lactate throughout anoxia and during at least the first day of aerobic recovery. Lactate is a glycolytic end-product that has classically been considered an unavoidable consequence of vertebrate anaerobic metabolism, but in WS40NE cells, 49 of 70 total unique carboxyethylation/lactylation modifications significantly changed in abundance in at least one comparison and one of these modifications was classified as highly condition dependent (Table 2). Given the extreme rarity of protein carboxyethylation, these modifications are most likely sites of lactylation [70]. Surprisingly, increased lactate accumulation in WS40NE cells is correlated with *decreased*, not increased, global carboxyethylation/lactylation. This pattern is the opposite of what is seen in human MCF-7, HeLa, and MDA-MB-231 cells, which exhibit a dose-dependent increase in histone lactylation in response to 25 mM exogenous L-lactate [69]. Further, in almost all cases, residues in WS40NE histones experience a decrease in lactylation during anoxia. It is possible that the slower rate of lactate accumulation in WS40NE cells (7.8 mM extracellular lactate after 4 days of anoxia) does not produce enough lactate to cause increase lactylation. However, lactate levels are relatively low during normoxia, and yet many histone residues still have a high percentage of lactylation. Thus, lactate availability is not likely to limit histone lactylation in WS40NE cells. It is worth considering that decreased rates of lactate accumulation and decreased histone lactylation in WS40NE cells may at least partially underlie their extreme anoxia tolerance, as this is contrary to trends that appear in anoxia intolerant mammalian cells.

### Dehydration and phosphorylation – a reciprocal relationship?

Dehydration of residues in WS40NE cell histones increases globally, while conversely, global phosphorylation decreases during anoxia and aerobic recovery from anoxia. These patterns are surprising, and the apparent reciprocal patterns of abundance are potentially very interesting.

One important site of phosphorylation was detected in WS40NE histones: H2A.X 137S. Phosphorylation of mammalian 139S (γ–H2A.X) initiates the DNA repair response and would therefore be expected to increase during anoxia due to an increase in ROS-induced DNA damage [71]. Surprisingly, 137S, the equivalent residue in WS40NE histone H2A.X, experiences a decrease in phosphorylation by 1 day in anoxia and continues to decrease during 4 days of anoxia. Interestingly, a nearby dehydrated residue, 142Y, follows the opposite pattern, increasing during anoxia. In this dataset, phosphorylation never occurs at this tyrosine residue. This is unexpected, as constitutive phosphorylation of mammalian 142Y by Williams–Beuren syndrome transcription factor (WSTF) occurs during normal conditions while dephosphorylation of this site occurs during the DNA damage response [72]. This dephosphorylation has been theorized to enhance recruitment of protein kinases to better maintain γ–H2A.X phosphorylation [72]. Therefore, it is interesting that this site is dehydrated, as dehydration may prevent phosphorylation and thus prevent the dampening of the DNA damage response that is observed in mammalian cells [72]. These sites should be explored in the future, as they are implicated in regulation of transcription and DNA replication may have nearby sites of dehydration that are involved in their modification status [73].

### Multiple hPTMs do not return to baseline after 1 day aerobic recovery

The histone landscape of WS40NE cells remains altered after 1 day of aerobic recovery from 4 days of anoxia. It is important to note that for these cells, 4 days is a relatively short duration of anoxia given they can survive for 49 days without oxygen [37]. Globally, 8 of the 13 types of hPTMs detected are significantly different in recovery compared to normoxia. Based on the log_2_ fold-change, global acetylation, oxidation/hydroxylation, and dioxidation are significantly elevated during recovery while global phosphorylation, biotinylation, 4-hydroxynonenal, methylation, and carboxyethylation/lactylation are all decreased. Of the unique hPTMs that were significantly modified, 42% are associated with recovery. While many of the hPTMs may simply not have returned to baseline, other modifications may be involved in recovery mechanisms. Further work will need to be done to determine if these hPTMs return to normoxic levels, or if they represent potential heritable epigenetic alterations to the histone landscape and act as a cellular memory of past exposures to anoxia.

### Conclusion

WS40NE cells are unique in their ability to survive long bouts of anoxia and demonstrate changes in their histone landscape in response to different lengths of time in anoxia and recovery from anoxia. This response includes altering both histone isoform relative abundance and hPTM relative abundance at multiple amino acid positions. This response likely contributes to the anoxia tolerance displayed in this cell type. Downstream effects of specific hPTM changes should be studied in the future to determine how specific hPTMs can impact gene expression and anoxia tolerance. Of specific interest is the potential reciprocal relationship between dehydration and phosphorylation as a regulatory mechanism to alter gene expression in response to anoxia.

## Methods

### WS40NE Cell Maintenance and Anoxic Exposure

Cells were grown in cell culture media (L-15 + 8.5% FBS, 5 mM glucose, 100 U/mL penicillin/ streptomycin) in 100 × 20 mm tissue culture dishes at 30°C and split when confluent using TrypLE Express [37]. For all anoxic experiments, plates of cells were seeded at 2.5 x 10^6^ cells per plate and given 24 h to adhere. Cells were then introduced into an anoxic chamber (Bactron EZ, Sheldon Manufacturing, Cornelius, OR, USA) that contains an atmosphere of 95% N_2_ and 5% H_2_. Media was immediately replaced with anoxic media that had been purged with N_2_ gas for 30 min and then pre-equilibrated to the atmosphere of the anoxic chamber overnight. Cells were harvested at **A)** 0 h of anoxia (normoxic), **B)** 1 d of anoxia, **C)** 4 d of anoxia, and **D)** after 1 d of aerobic recovery following 4 d of anoxia (n = 12). To harvest, media was removed and a cell scraper was used to gently dissociate cells from the bottom of the plate. Two 100 × 20 mm tissue culture dishes were pooled for each replicate. Cell pellets were generated by subjecting the isolated cells to centrifugation at 500 x *g* for 7 min at 4° C, and removing the supernatant. Additionally, media samples were taken directly prior to the harvest of all four timepoints and used to determine the levels of extracellular lactate accumulation. Cell pellets and spent media were flash frozen in liquid nitrogen. Samples were stored at -80° C until use for histone acid extraction or determination of lactate concentration.

### Histone Acid Extraction and Protein Digestion

A middle-down approach to proteomics was used for liquid chromatography mass spectrometry (LCMS) using previously described methods [34, 43]. This workflow was chosen due to its previous success in capturing histone peptide fragments in tissue of adult Mozambique tilapia (*Oreochromis mossambicus*) and *A. limnaeus* embryos [34, 43]. Histone acid extracts were generated by resuspending cell pellets in 0.4 N H_2_SO_4_ and incubating the samples with constant rotation at 4 °C for 4 h. Samples were then centrifuged at 16,000 x *g* for 10 min at 4 °C and the supernatant was transferred into a fresh 1.5 mL tube. This process was repeated twice. Trichloroacetic acid (TCA) was added to each sample drop-by-drop until a volume equal to 25% of the estimated sample volume had been added. Samples were inverted to mix and incubated overnight at 4 °C. Once the histone proteins had precipitated, samples were centrifuged at 3,400 x *g* for 5 min at 4 °C, and the supernatant was discarded. The remaining pellet was rinsed twice: first in ice cold acetone with 0.1% hydrochloric acid and then with 100% acetone. For both rinses, the samples were centrifuged at 3,400 x *g* for 2 min at 4 °C and the supernatant was removed. After both rinses were complete, histone pellets were air dried in a fume hood for 5 min. Samples were stored at -80 °C until the in-solution digestion.

To begin the digestion, each sample was brought to room temperature (RT), and the histone pellet was resuspended in 138 μL of 8 M urea. Dithiothreitol (DTT) was added to a final concentration of 10 mM. Each sample was then briefly vortexed, centrifuged, and incubated at 37 °C for 30 min. Iodoacetamide (IAA) was added to each sample to a final concentration of 30 mM, and samples were briefly vortexed and centrifuged before a 30 min incubation in the dark at RT. Samples were briefly vortexed and centrifuged again after the incubation and protein concentration was determined through a bicinchoninic acid (BCA) assay, (cat# 23250, Thermo Scientific, Waltham, MA, USA). Each sample (50 μg of total protein) was diluted 5:6 (v:v) in 0.1M sodium phosphate buffer (pH 7.8). V8 protease beads (cat#20151, Thermo Scientific, Waltham, MA, USA) were added at 1:50 ratio (protease:total protein). In sodium phosphate buffer, V8 protease cleaves proteins at the carboxyl end of both glutamate and aspartate residues. After a 20 h incubation with constant rotation at 30 °C, protease beads were removed by centrifugation at 500 x *g* for 2 min and the supernatants were retained. Supernatants were subjected to an additional centrifugation at 19,000 x *g* for 5 min, and those subsequent supernatants were transferred to fresh tubes and dried using a SpeedVac (Thermo Savant, model# ISS110, Thermo Scientific, Waltham, MA, USA). The peptide pellets were reconstituted to a final concentration of 333 ng/μL in 0.1% formic acid diluted in LCMS-grade water. For peptide clean-up, Pierce C18 Spin Columns (cat# 89873, Thermo Scientific, Waltham, MA, USA) were used according to the manufacturer’s instructions.

Data-dependent acquisition (DDA) was used to generate peak lists [74], and PEAKS Suite X Plus and MSFragger were used to annotate histone peptides using the *A. limnaeus* reference proteome [75]. Utilizing the spectral library that was generated from the DDA run, data-independent acquisition (DIA) was then used to quantify the peptides in the samples using fragment ion–based extraction. A fixed isolation window DIA scheme was applied, with consecutive 10 m/z windows overlapping by 0.5 m/z. The full MS/MS acquisition range spanned *m/z* 370–1090 and collision energy was optimized to specifically match each isolation window. For each peptide, at least four transition peaks were required to be included in the dataset [43, 74]. PTMs were assigned by PEAKSPTM software using a minimum localization score of 5% and a minimum Ascore of 20. All data were normalized in Skyline using overall sample median.

### Histone and hPTM Library Generation

Peptides modified with hPTMs were distinguished from unmodified sequences based on their mass shift (Fig. S1). A list of histone PTMs was compiled according to the histone protein name, amino acid residue/position, and modification type using Python code as previously described [34]. All histone PTMs detected were evaluated for their biological relevance (classified as post-translational, other, or multiple) according to their Unimod accession and amino acid residue. Modifications classified by Unimod as known artefacts or chemical derivatives were removed from analysis. Modifications with the same mass shift are listed together. Peptides that mapped to multiple histones in the *A. limnaeus* genome are presented as a protein family and are reported together.

**Figure S1:**
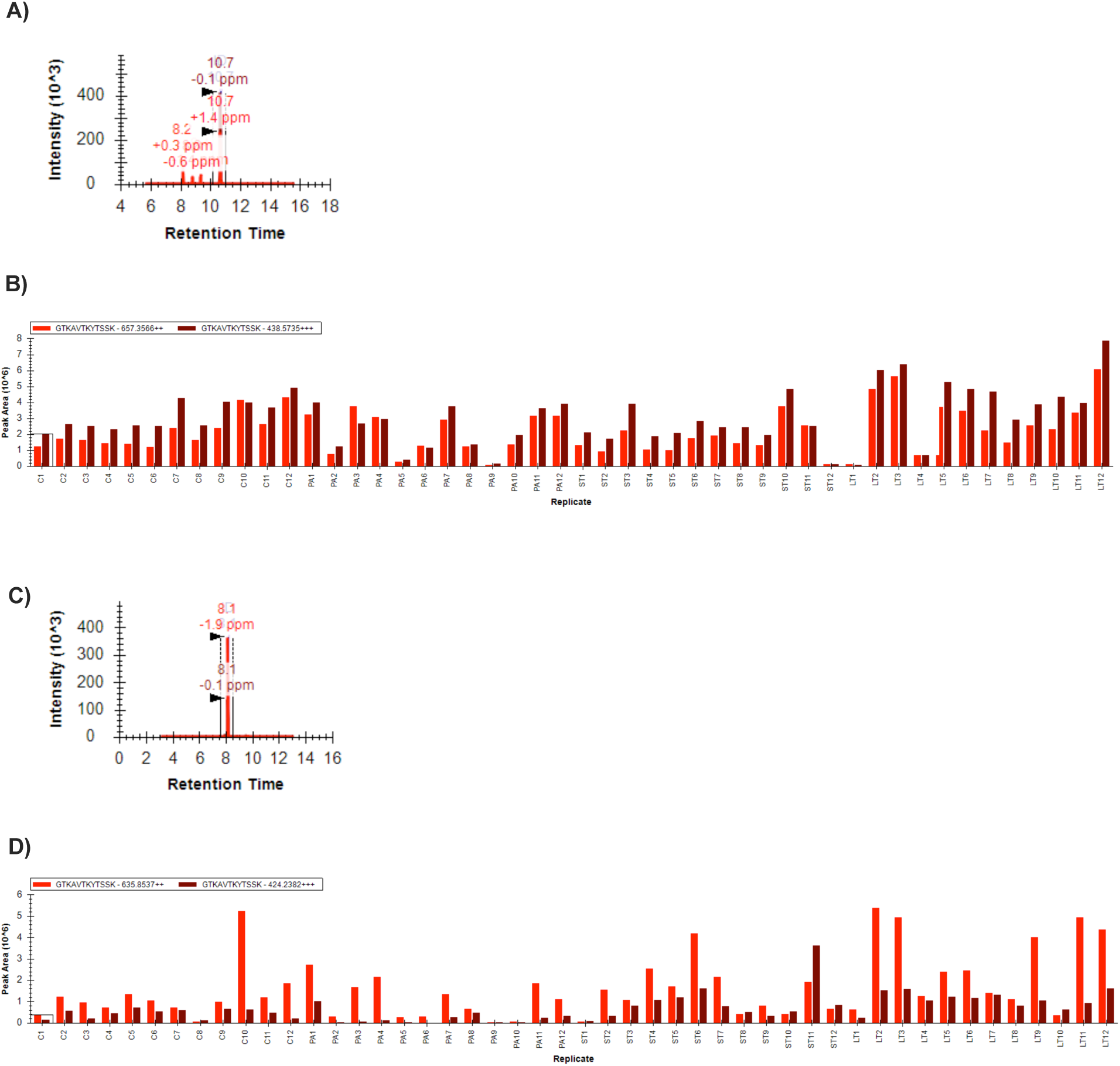
Quantification of histone post-translational modifications in Skyline. A representative MS spectrum (**A**) and peak area (**B**) of the most abundant ions for a modified peptide, E.GTK[+42]AVTKYTSSK, and **(C,D)** the corresponding unmodified peptide E.GTKAVTKYTSSK. Both peptides correspond to histone H2B.L4-like (XP_013885354.1). A mass shift of +42 on a peptide represents acetylation.

### Protein Abundance and hPTM Statistical Analyses

Histone isoform abundance was evaluated using quantile normalized raw protein abundance data in Python as previously described [34]. Principal component analysis (PCA) and two-tailed t-tests with the Benjamini-Hochberg correction applied to account for multiple hypothesis testing [76] were used to make inferences about histone isoform abundance.

Peptide abundance, as determined by normalized area under the curve, and raw protein abundance were obtained using Skyline (Version 22.2) [77]. Relative abundance values (the percent that a specific amino acid residue on a histone is modified by a specific PTM) were used to calculate beta-values and M-values. Beta-value is a transformation of relative abundance expressed as a number between 0 and 1 calculated by adding a value of 100 to the denominator to standardize low intensity peptides [78]. The M-value is a logit transformation of the beta-value [79]. Additionally, the relative abundance of each hPTM was used to calculate the log_2_ fold change, and a two-tailed t-test was used to compare the relative abundance of hPTMs in all conditions with the Benjamini-Hochberg correction applied to account for multiple hypothesis testing [76]. This relative abundance analysis was repeated to analyze global hPTM levels and calculate the log_2_ fold change. Additionally, an ANOVA was performed on the global relative abundance of each hPTM using Prism (ver 10.0.0, GraphPad, Software, Boston, MA, USA). A Dunnett’s multiple comparison test was used to determine significance of all conditions compared to normoxia. The global relative coverage of each modifiable amino acid residue was calculated to determine what percentage of each hPTM was present at each modifiable residue.

Histone PTM maps were constructed for anoxic and normoxic samples using the mean relative abundance of hPTMs. The M-value was used to perform a principal component analysis (PCA). All abundance plots and PCA plots were made in R using the package ggplot2 [80]. Global hPTM abundance, protein relative abundance, and global relative residue coverage were visualized using Prism (ver 10.0.0, GraphPad, Software, Boston, MA, USA). All Python and R code is accessible by the public on GitHub at https://github.com/hughcj11/WS40NE_Histone_PTM_Analysis. Supplementary data including average relative abundance, t-statistic, p-value, and corrected p-value is provided for each unique, biologically relevant hPTM are provided in Supplementary Table S1 (Excel workbook) across all aerobic conditions.

### Lactate Accumulation Assay

Lactate concentration in the cell culture media was measured enzymatically as previously described [37]. Briefly, each well of a 96-well plate (black walls, clear bottom, Greiner Bio-One, Monroe, NC, USA) contained the following enzyme assay cocktail: 218 μL of Glycine (0.6 M)-Hydrazine (0.5 M) buffer (pH = 9.2), 25 μL 27 mM NAD^+^, 5 μL of lactate standard or sample, and 2.5 μL lactate dehydrogenase (LDH, 14 U/mL, Sigma Aldrich). Change in absorbance was measured at 340 nm every min for 60 min using a plate reader (Tecan Infinity M200Pro, Männendorf, Switzerland). Final concentration of lactate in the samples was determined by interpolation using a lactate standard curve.

## Author Contributions

JEP and CH conceived and designed the study. CH and EAM performed the experiments and acquired the data. CH analyzed the data. DK and JEP contributed reagents/materials/analysis tools. CH and JEP wrote the paper. All authors edited and approved the manuscript.

## Acknowledgements

We would like to thank Karim Soliman for his guidance in the creation of the Python data analysis pipeline and visualization of the data. We would like to thank Dr. Daniel Zajic for his help performing the lactate accumulation assay.

## Data and resource availability

The datasets (DDA and DIA raw data) supporting the conclusions of this article are available in Panorama Public (https://panoramaweb.org/CH0001JP.url) and ProteomeXchange (PXD067707: https://proteomecentral.proteomexchange.org/cgi/GetDataset?ID=PXD067707).

Reviewer login:

Password: 3dqk$TMm1AJbo%

